# Neural Network-Assisted Humanization of COVID-19 Hamster scRNAseq Data Reveals Matching Severity States in Human Disease

**DOI:** 10.1101/2024.01.11.574849

**Authors:** Vincent D Friedrich, Peter Pennitz, Emanuel Wyler, Julia M Adler, Dylan Postmus, Luiz Gustavo Teixeira Alves, Julia Prigann, Fabian Pott, Daria Vladimirova, Thomas Hoefler, Cengiz Goekeri, Markus Landthaler, Christine Goffinet, Antoine-Emmanuel Saliba, Markus Scholz, Martin Witzenrath, Jakob Trimpert, Holger Kirsten, Geraldine Nouailles

**Affiliations:** University of Leipzig, Institute for Medical Informatics, Statistics, and Epidemiology, Leipzig, Germany; Center for Scalable Data Analytics and Artificial Intelligence (ScaDS.AI), Leipzig, Germany; Charité – Universitätsmedizin Berlin, corporate member of Freie Universität Berlin and Humboldt-Universität zu Berlin, Department of Infectious Diseases, Respiratory Medicine and Critical Care, Berlin, Germany; Max Delbrück Center for Molecular Medicine in the Helmholtz Association (MDC), Berlin Institute for Medical Systems Biology (BIMSB), Berlin, Germany; Freie Universität Berlin, Institut für Virologie, Berlin, Germany; Charité – Universitätsmedizin Berlin, corporate member of Freie Universität Berlin and Humboldt-Universität zu Berlin, Institute of Virology, Berlin, Germany; Berlin Institute of Health at Charité – Universitätsmedizin Berlin, Berlin, Germany; Liverpool School of Tropical Medicine, Department of Tropical Disease Biology, Liverpool, United Kingdom; Gladstone Institutes, San Francisco, USA; Cyprus International University, Faculty of Medicine, Nicosia, Cyprus; Humboldt-Universität zu Berlin, Integrative Research Institute for the Life Sciences, Berlin, Germany; Faculty of Medicine, Institute of Molecular Infection Biology (IMIB), University of Würzburg, Würzburg, Germany; Helmholtz Institute for RNA-based Infection Research (HIRI), Helmholtz Center for Infection Research (HZI), Würzburg, Germany; University of Leipzig, Faculty of Mathematics and Computer Science, Leipzig, Germany; German Center for Lung Research (DZL), Berlin, Germany

**Author notes:** Corresponding Authors: Holger Kirsten, Dr. rer. nat., Institute for Medical Informatics, Statistics, and Epidemiology (IMISE), Haertelstraße 16-18, 04107 Leipzig, Germany, Geraldine Nouailles, Dr.-Ing., Charité – Universitätsmedizin Berlin, Corporate Member of Freie Universität Berlin and Humboldt-Universität zu Berlin, Department of Infectious Diseases, Respiratory Medicine and Critical Care, Charitéplatz 1, 10117 Berlin, Germany, / /. co-first (Vincent D Friedrich, Peter Pennitz). co-last authors (Holger Kirsten, Geraldine Nouailles).

## Abstract

Translating findings from animal models to human disease is essential for dissecting disease mechanisms, developing and testing precise therapeutic strategies. The coronavirus disease 2019 (COVID-19) pandemic has highlighted this need, particularly for models showing disease severity-dependent immune responses. Single-cell transcriptomics (scRNAseq) is well poised to reveal similarities and differences between species at the molecular and cellular level with unprecedented resolution. However, computational methods enabling detailed matching are still scarce. Here, we provide a structured scRNAseq-based approach that we applied to scRNAseq from blood leukocytes originating from humans and hamsters affected with moderate or severe COVID-19. Integration of COVID-19 patient data with two hamster models that develop moderate (Syrian hamster, *Mesocricetus auratus*) or severe (Roborovski hamster, *Phodopus roborovskii*) disease revealed that most cellular states are shared across species. A neural network-based analysis using variational autoencoders quantified the overall transcriptomic similarity across species and severity levels, showing highest similarity between neutrophils of Roborovski hamsters and severe COVID-19 patients, while Syrian hamsters better matched patients with moderate disease, particularly in classical monocytes. We further used transcriptome-wide differential expression analysis to identify which disease stages and cell types display strongest transcriptional changes. Consistently, hamster’s response to COVID-19 was most similar to humans in monocytes and neutrophils. Disease-linked pathways found in all species specifically related to interferon response or inhibition of viral replication. Analysis of candidate genes and signatures supported the results. Our structured neural network-supported workflow could be applied to other diseases, allowing better identification of suitable animal models with similar pathomechanisms across species.

**Key Points:**

- Neural networks can successfully match disease states between animal models and humans using single-cell data as shown for COVID-19
- Moderately diseased patients best matched Syrian hamster cells; severely diseased patients best matched Roborovski hamster neutrophils

## Introduction

Animal models are essential for understanding host-pathogen interactions^1^ and for evaluating new therapies, as demonstrated by the rapid response to the recent severe acute respiratory syndrome coronavirus type 2 (SARS-CoV-2) caused pandemic, which was greatly aided by animal studies^2-4^. Nonetheless, the ‘translational gap’ from animal models to human disease often remains a major challenge^5^. One means to minimize this gap is to implement unbiased tools, i.e. tools not requiring explicit biological pathway knowledge, such as neural network-based models.

COVID-19 presents a suitable disease for conducting proof-of-concept studies. Multiple well-documented and publicly accessible single-cell transcriptomic (scRNASeq) datasets derived from blood of COVID-19 patients are available^6-10^. COVID-19 patients exhibit a range of disease severities, which is measured using the world health organization’s (WHO) ordinal scale for disease severity^11^ (WOS). Various suitable COVID-19 animal models, such as non-human primates, ferrets, hamsters, and transgenic mice, typically addressing specific human disease stages, have been reported^12-16^. Hamster models are particularly suitable, as Syrian hamsters (*Mesocricetus auratus*) develop a moderate and Roborovski hamsters (*Phodopus roborovskii*) a severe disease course upon experimental infection^17-19^. However a comprehensive cross species comparison especially on the molecular level is still an unmet need.

For an unbiased and omics-based comparative analysis across species, individual species datasets first need to be successfully integrated. Multiple approaches exist, with different trade-offs between eliminating batch effects and preserving biological variance^20^. Integration of information across multiple experiments was demonstrated by the so-called Human Lung Cell Atlas (HLCA) core data set^21^, where a neural network-based method called scANVI^22^ was used to integrate 14 datasets.

Neural network-based models can also assist in comparing biological signals and patterns in high-dimensional scRNAseq data. Variational Autoencoders (VAEs) are especially valuable for this purpose^23,24^. VAE models can represent gene expression data (about 20,000 genes) in a low-dimensional space using as few as 10 dimensions, while also capturing the most significant input signals. Thus, VAEs have been utilized in scRNAseq data analyses to model sparse data^25^, integrate data^26,27^, generate data^27-29^ and perform reference mapping^30^. The development of scGen by the lab of Theis and colleagues is a vital application of VAEs^27^. scGen has the capability to deduce data points for conditions not present in the data. It successfully predicted LPS responses across various animal species^27^, as well as protein abundance from mRNA in cases of Alzheimer’s dementia^31^, and gene expression patterns in mice, humans, and cancer cell-lines following chemical perturbations^32^.

Here, we employs an scGen-based framework to match COVID-19 disease severity levels across humans and hamster species, while also developing a metric for measuring interspecies similarity on a cellular level. To complement this, we used differential expression analysis to compare up- and downregulation at whole genome, candidate-gene and pathway level, across different species and severities. Our study demonstrates how analysis of omics datasets across different species, utilizing VAEs and other bioinformatics methods, can help to identify strengths and limitations of specific animal models and their relationships to human disease states.

## Materials and Methods

### Ethics statement and blood sampling for scRNAseq analysis

All experiments involving animals were approved by relevant institutional and governmental authorities (Freie Universität Berlin and Landesamt für Gesundheit und Soziales Berlin, Germany, permit number 0086/20) and adhered to the Federation of European Laboratory Animal Science Associations (FELASA) guidelines and recommendations for the care and use of laboratory animals, which is equivalent to American ARRIVE guidelines. Detailed description of experimental animals, virus stock and infection procedure can be found in **Supplemental Methods**. Briefly, 500 µL whole blood per animal was depleted of red blood cells by lysis, counted and processed for single-cell RNA sequencing using 3’-based 10x Genomics chemistry according to the manufacturer’s instructions and as we previously published^33^. Sequencing was performed on a Novaseq 6000 device (Illumina) according to the manufacturer’s instructions resulting in sample averages of 34,387 reads per cell and sequencing saturation of 73.1% (**Supplementary Table1**). Further information regarding cell processing is available in **Supplemental Methods**.

### Data integration

Orthologue genes were defined based on Ensembl release 109 - Feb 2023, for *Phodopus roborovskii,* orthologue information from *Mesocricetus auratus, Rattus_norvegicus, or Mus_musculus* was used as available (**Figure S1, Supplementary Table2**). The datasets of the three species (including cells marked as low quality) were integrated using anchor-based reversed PCA as implemented in the IntegrateLayers() function of the R-package Seurat. Cell clusters were identified again and visualized by embedding expression profiles in a Uniform Manifold Approximation and Projection (UMAP). For each cluster of cells, we identified the most likely cell type according to the originally reported cell label which was in line with reported marker genes. Cells with discordance between cluster and original label were removed (5.5%). Clusters from single species and integrated analyses with more than 50% cells not meeting cut-offs were removed if not explained by cell type (i.e. neutrophils and plasmablast cluster). Individual low-quality cells were also removed, in total 6.3% of all cells (**Supplementary Table3**).

### VAE disease state matching

We applied variational autoencoders (VAEs) for jointly embedding high-dimensional hamster and human gene expression data into a low-dimensional latent space using the python package scGen^27^ v2.0.0. Separate VAE models were trained for comparisons between humans and Syrian hamsters as well as between humans and Roborovski hamsters. To account for diverse transcriptomic responses to infection across cell types, distinct VAE models were learned for each cell type. General interspecies differences in latent space were addressed by a species-shift-vector based exclusively on human controls (WOS0) and hamster controls (Day0). The control group-based species-shift-vector ensured controlling for interspecies difference while avoiding interference from infection response. To allow for cross-species disease state comparison, the species-shift-vector derived from the controls was applied to the latent embeddings of infectious hamster cells, a process we refer to as humanization. The similarity between humanized-hamster disease states and human disease states was quantified via a diffusion pseudotime-based^34^ distance metric *d2*. Details can be found in **Supplemental Methods**.

### Statistical Analysis

R-package FDRtool 1.2.17 was used to estimate percentages of differential transcriptome in cells of infected vs. healthy subjects. Differential expression analysis of time points after infection compared to controls in hamsters, and WOS levels compared to controls in humans, were performed in the Limma-Voom^35^ framework, standard-filtering for frequent genes using pseudo counts aggregated per biological replicate. Multiple testing of multiple genes was controlled by the false-discovery-rate according to Benjamini and Hochberg. To correlate fold-changes of genes expressed in the same direction in human patients and one hamster species, all genes significant at the level of FDR≤20% in both species were used. When comparing the top-10 differential regulated genes “corresponding direction” means either the same direction or, for timepoint 14 days post infection (dpi; i.e. recovered hamsters), the opposite direction between patients and hamsters. Pathway enrichment analyses were performed using gprofiler2 0.2.1. filtering the databases Biological Processes from Gene Ontology, Wikipathways, Reactome, and KEGG. In this way, the foreground was defined as maximum 100 top-genes at FDR≤20% ordered by absolute fold-change or - if this resulted in less than 30 genes - the top 30 genes ordered by *P*-value, excluding ribosomal and mitochondrial genes. The background were all genes considered in association analysis. When comparing overlapping pathways between species, we filtered for pathways being globally significantly enriched (padjusted ≤0.05) and including at least 2 foreground-genes.

### Data Availability

Publicly available datasets that were used in this manuscript can be found at GEO:“GSE162208” (Syrian hamsters) and EGA:“EGAS00001004571” (humans). Datasets for Roborovski hamsters and processed data will be available via zenodo and GEO. The code used for data analysis is available at github.com, https://github.com/GenStatLeipzig/Neural-Networks-Assisted-Humanization-of-COVID-19-Hamster-scRNASeq-data.

## Results

We generated scRNAseq from blood of uninfected or infected Roborovski hamster (n=3 per time point and condition) that received a low-dose (1 × 10^4^ pfu) or a high-dose (1 × 10^5^ pfu) of SARS-CoV-2 to compare with our previously published data from infected (1 × 10^5^ pfu) and uninfected Syrian hamsters^33^ and publicly available human data from COVID-19 patients and controls^7^ (**Figure 1A**). All together the data encompass 28357 cells in Syrian hamsters, 23046 cells in Roborovski hamsters and 79196 cells from humans.

**Figure 1.**
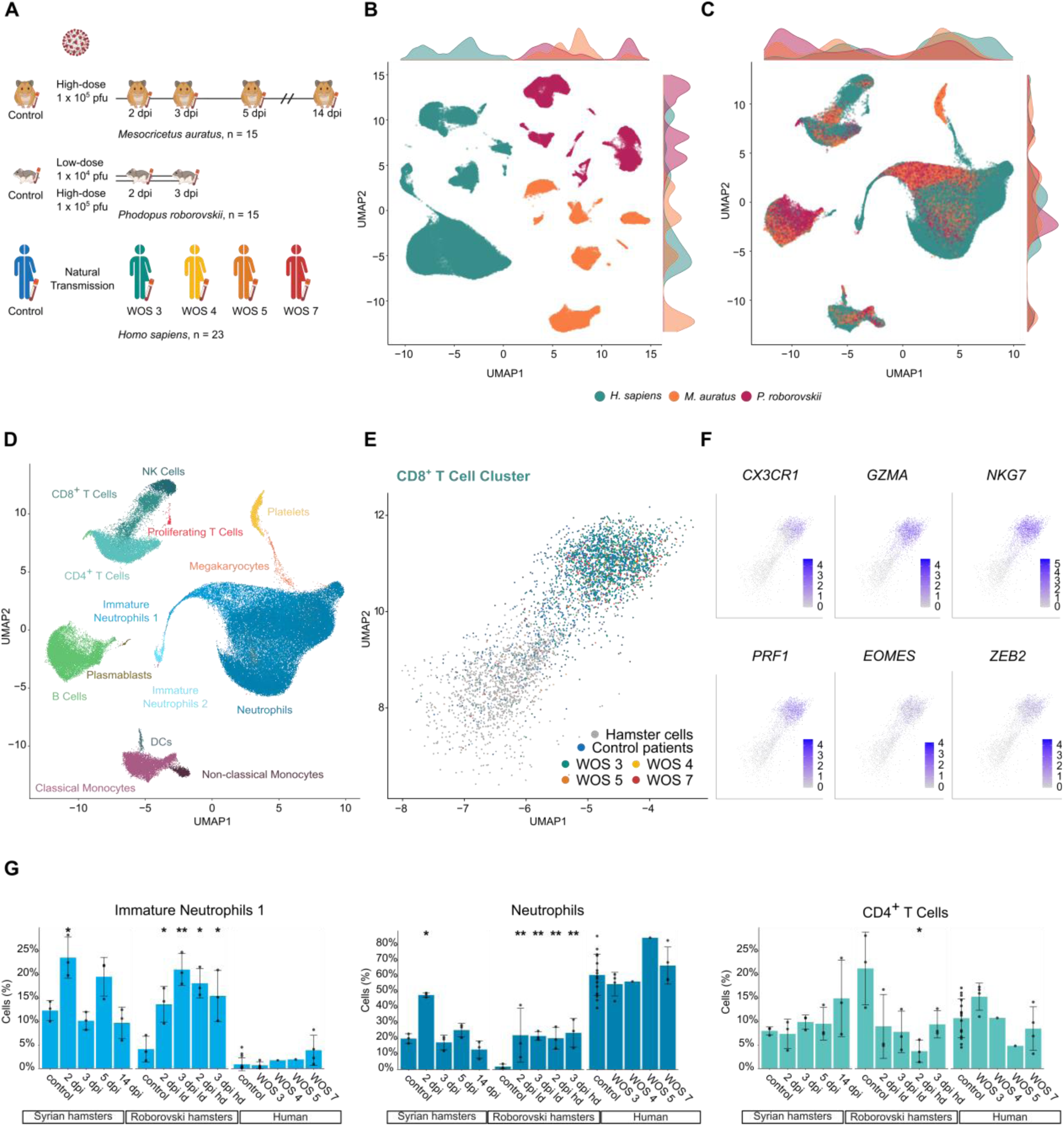
Cellular recruitment in hamsters and humans. (A) Schematic overview of included datasets from humans and hamsters. (B) Uniform manifold approximation and projection (UMAP) plot indicating the original dataset of all cells merged in a common embedding before data integration including cell density per UMAP coordinate colored according to species. (C) UMAP plot indicating the original dataset of all cells after performing data integration using Reciprocal Principal Component Analysis (RPCA) including cell density per UMAP coordinate colored according to species. (D) UMAP plot of identified cell populations of all datasets. (E) UMAP Plot of CD8^+^ T cell subset of human and hamster cells; colors indicate cells from different human severity levels (F) UMAP plots of CD8^+^ T cell subset indicating the expression of genes linked to cytotoxicity, and transcription factors associated with T cell effector/memory differentiation. (G) Frequencies of different populations in the two hamster species according to days post-infection and in humans according to the WHO ordinal scale for disease severity (WOS). To quantify significance of differential abundance of cell types, we applied a negative binomial generalized linear model as previously described^58^. * FDR ≤ 0.05, ** FDR ≤ 0.01. FDR: global false discovery rate. Data display means ± SD. n = 3 animals per time point for hamsters. Human Data: Cohort 2 from Schulte-Schrepping, *et al.^7^* (**Supplemental Table9**).

### Cellular recruitment in hamsters and humans

First, we combined blood transcriptomic data into a common embedding and applied a single unified gene nomenclature as described previously^36^. The merged dataset revealed species-related batch effects (**Figure 1B**). To allow for application of a common cell type assignment, we resolved these by utilizing the Reciprocal Principal Component Analysis (RPCA) data integration pipeline^37^ (**Figure 1C**). Application of Louvain-based clustering to integrated interspecies data identified several matching leukocyte populations (**Figure 1D**). One previously described^7^ subset of immature neutrophils, namely *FUT4*^+^ pro-neutrophils (Immature Neutrophils 2), was exclusively observed in humans. In addition, while CD8^+^-T-cells from hamsters clustered closer to CD4^+^-T-cells, CD8^+^-T-cells from COVID-19 patients clustered closer to NK-cells (**Figure 1E**), involving genes linked to cytotoxicity (*GZMA*, *NKG7*, *SLAMF7*, *CX3CR1*, *PRF1*), as well as transcription factors associated with T-cell effector/memory differentiation (*ZEB2*, *EOMES*) (**Figure 1F**).

Examining the proportion of blood cell types over time after infection in hamster data, or with increasing disease severity in human data, revealed distinct patterns: Syrian hamsters experienced a sharp but transient increase in blood neutrophils at 2 days post-infection (dpi), resolving to basal levels by 3dpi. In contrast, in both high- and low-dose infected Roborovski hamsters, neutrophils remained increased at and 3dpi. In humans, a steady increase in the immature neutrophil fraction correlated with disease severity. The fraction of CD4^+^-T-cells declined in patients with severe disease (WOS5, WOS7) compared to controls, which was consistent with both Roborovski hamster infection groups. By contrast, in patients with moderate disease course (WOS3, WOS4), CD4^+^-T-cells increased. A reduction of non-classical monocytes, which is common in severe human disease^6,7^, was not observed in infected hamsters (**Figure 1G, S2**).

### VAE-based neural network pipeline enables interspecies disease state matching of the transcriptome

To identify transcriptional similarities between humans and hamsters during early stages of SARS-CoV-2 infection, we employed neural network techniques to match disease states across species. High-dimensional gene expression data was mapped to low-dimensional latent space through implementation of a variational autoencoder (VAE) model^27^. Latent embedding provides a denoised data representation, capturing only the most relevant input signals (**Figure 2A**). For each hamster species and every cell type of interest, we trained separate VAE models for jointly embedding human and hamster data.

**Figure 2.**
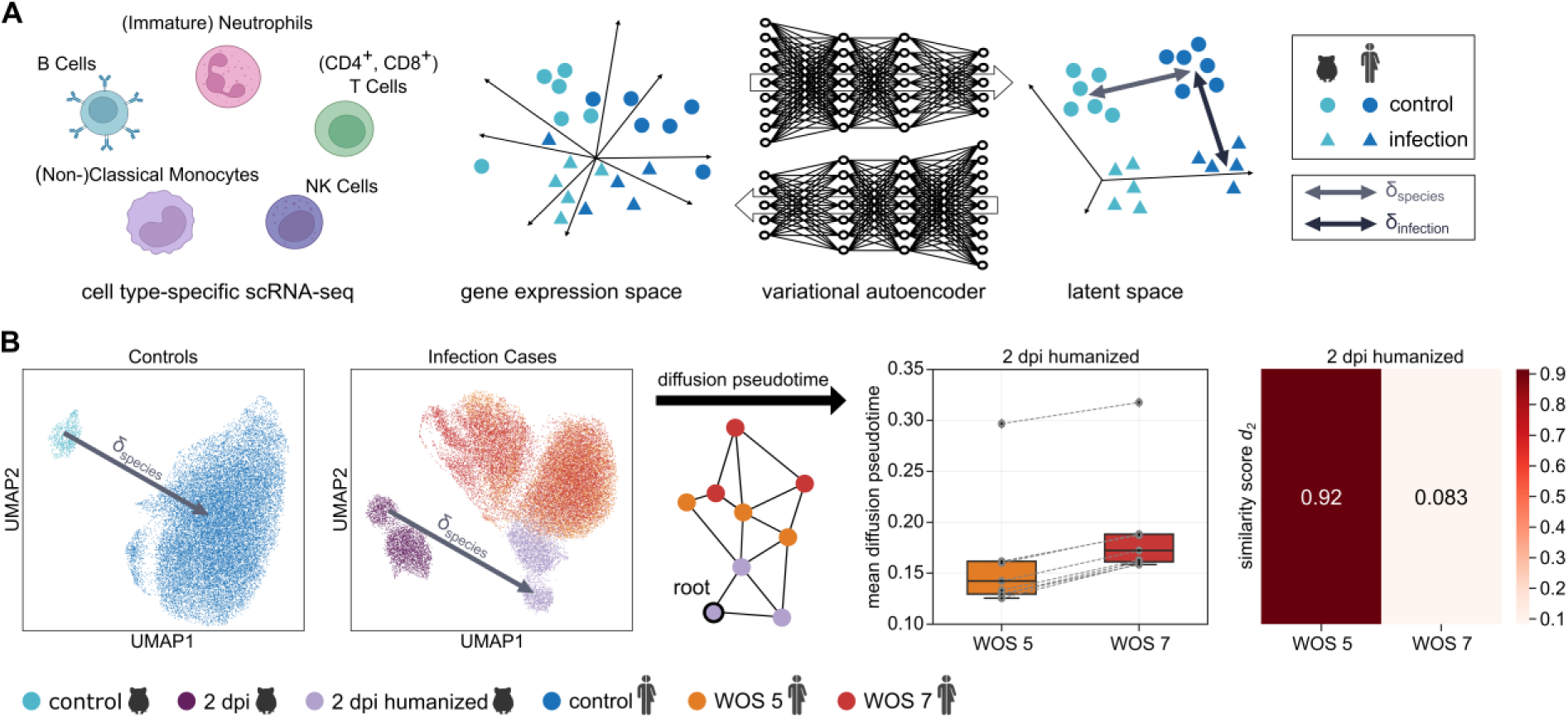
VAE neural network-based workflow enables interspecies disease state matching of the transcriptome. (A) Schematic overview of the variational autoencoder (VAE) neural network-based pipeline for joint latent embeddings of high-dimensional hamster and human cell type-specific scRNA-seq data. (B) Disease state matching in VAE latent space illustrated for Syrian hamster and human neutrophils with selected disease states. The species-shift-vector *δspecies* is derived from Syrian hamster and human controls (left panel, UMAP coordinates adapted) and applied to humanize the latent embedding of 2 dpi Syrian hamster cells (second left panel, UMAP coordinates adapted). To identify the best matching WOS grade, the mean diffusion pseudotime distances (graph-based, scheme middle panel) between the 2 dpi humanized Syrian hamster disease state and the human disease state WOS 5 are calculated, as well as with the WOS 7 state, respectively. For this we used 10 different root cells (second right panel), converted this information into a similarity score *d2* (right panel) and identified the best matching disease state.

Figure 2B illustrates the procedure to match disease states of human and Syrian hamster neutrophils using the example of finding the human severity grade corresponding best with Syrian hamster 2dpi. In latent space, the species-shift-vector (*δspecies*) is derived based on uninfected human and hamster controls via vector arithmetic (**Supplemental Methods**). Since only control groups were considered for computing *δspecies,* it represents species differences, but not response to infection. The humanized representation of each hamster cell 2dpi in latent space was then generated by applying *δspecies* to the cell’s latent embedding and compared to the latent embeddings of human WOS5 and WOS7 cells by a similarity quantification of species disease states using graph-based diffusion pseudotime distance^34^ incorporating 10 different root cells. For all root cells, humanized-hamster cells 2dpi exhibited smaller mean diffusion pseudotime distance to human WOS5 cells (similarity score *d2* = 0.92) than to WOS7 cells (*d2* = 0.083) (Figure 2B).

### Species- and infection-shift-vectors in VAE latent space generalize to out-of-training samples

To evaluate the generalizability of the *δspecies* approach, we divided human and Syrian hamster neutrophil control groups into a training set and a test set (Figure 3A, upper panel). *δspecies* was derived *de novo* solely from the VAE latent embeddings of the training set. Next, we applied the VAE encoder, *δspecies* and VAE decoder from the training set to the test set data, resulting in a decoded test set (**Supplemental Methods**). The application was considered successful, since the decoded test set of humanized-hamster cells displayed a stronger correlation with human data compared to the original decoded hamster data. Correlation was quantified using squared Spearman’s rank correlation coefficient *R^2^* of all gene-wise calculated medians of species data with values ranging between 0 (no correlation) and 1 (perfect correlation). Decoded humanized-hamster achieved a higher *R^2^* score (0.97) compared to decoded original hamster (0.11) when tested against the gene expression of the human control group (Figure 3A, lower panel) demonstrating generalizability of *δspecies* to out-of-training samples. Next, confirmation that humanization works for both hamster species was achieved by repeating the training-test-split humanization procedure with non-classical monocytes from Roborovski hamsters - again resulting in a higher *R^2^* score for decoded humanized-hamster (0.84) compared to decoded original hamster (0.27, Figure 3B). Analogously, we tested the remaining cell types of both hamster species; humanization worked properly at all occasions with best results for Syrian hamster (**Figure S3**).

**Figure 3.**
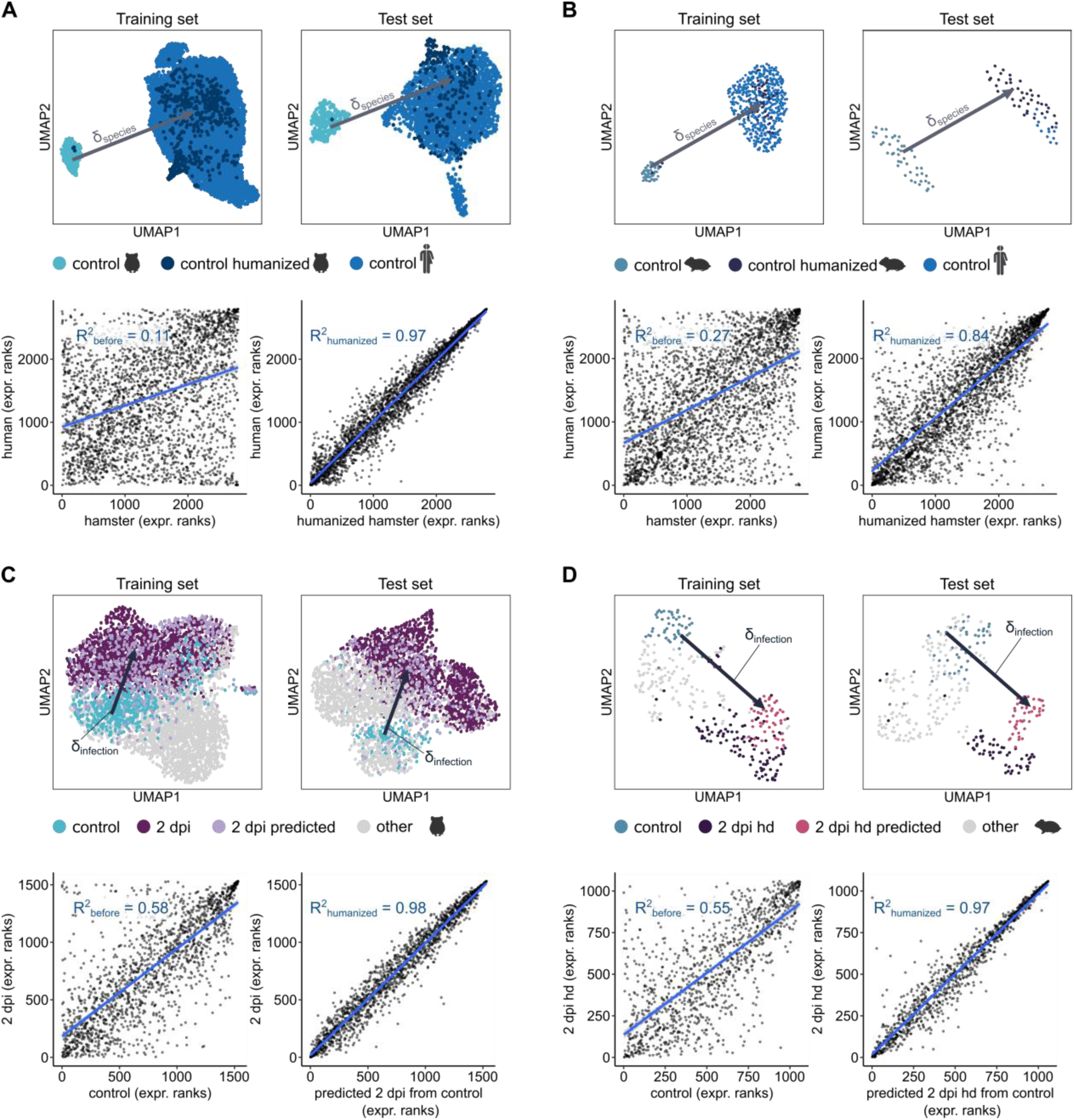
Species- and infection-shift-vectors in VAE latent space generalize to out-of-training samples. (A) Cross-species proof-of-principle for Syrian hamster and human neutrophil controls: The species-shift-vector *δspecies* in VAE latent space is derived solely from the training set (top left panel, UMAP coordinates adapted) and is applied to humanize the latent embeddings of hamster control cells of the test set (top right panel, UMAP coordinates adapted). Rank correlation scatter plots for decoded hamster and human gene expression (bottom left panel) as well as for decoded humanized hamster and human gene expression (bottom right panel) of the test set. (B) Analogous to (A) for Roborovski hamster and human non-classical monocyte controls. (C) Intra-species proof-of-principle for Syrian hamster neutrophils: The infection-shift-vector *δinfection* is derived solely from the training set and points from control group towards the 2 dpi disease state in VAE latent space (top left panel, UMAP coordinates adapted). The 2 dpi disease state of control cells from the test set is predicted by applying the infection-shift-vector *δinfection* from the training set in VAE latent space (top right panel). Rank correlation scatter plots for decoded control and 2 dpi gene expression (bottom left panel) as well as for decoded predicted 2 dpi and control gene expression (bottom right panel) on the test set. (D) Analogous to (C) for Roborovski hamster non-classical monocyte controls with predicted high-dose 2 dpi states of the control group. *R^2^*: Spearman’s rank correlation coefficient.

We repeated the described training-test-split procedure to evaluate robustness of infection-shift-vectors *δinfection* representing the mean direction of infection in latent space and enabling prediction of diseased states from a control cell. For the example of Syrian hamster neutrophils 2dpi, the decoded predicted gene expression for 2dpi using control cells not used for training correlated more strongly with the decoded actual gene expression at 2dpi (*R^2^*: 0.98) than with the corresponding control cell gene expression (*R^2^*: 0.58) on the test set (Figure 3C). Similar results were found for Roborovski hamster non-classical monocytes (*R^2^*: 0.97 vs. *R^2^*: 0.55, respectively) (Figure 3D). This demonstrates that disease states can be addressed by an infection-shift-vector *δinfection* in VAE latent space, which generalizes to out-of-training samples.

### VAE-driven mapping of hamster temporal disease states to patient severity levels

The VAE model pipeline was then used to map temporal disease states in hamsters to levels of disease severity in COVID-19 patients across all datasets and cell types with sufficient cell counts. At all time points examined, resulting transcriptional states of most Syrian hamster cell types best matched patients with moderate disease based on similarity score *d2*, particularly WOS3 and 4 (Figure 4A**, S4A, S4B**). However, for some cell types no clear assignment to a particular human severity grade could be established. For example, transcriptomic profiles of Syrian hamster non-classical monocytes corresponded not only to that of WOS3 patients, as expected, but also resembled that of patients with WOS7. Among the Roborovski hamsters’ blood leukocytes (Figure 4B**, S4C, S4D**), neutrophils, independently of infection dose, demonstrated highest similarity to WOS7 patients. Conversely, classical monocytes matched transcriptomic profiles of patients with moderate disease courses. Immature neutrophils of both hamster species matched moderate patients but also showed similarity to severe patients.

**Figure 4.**
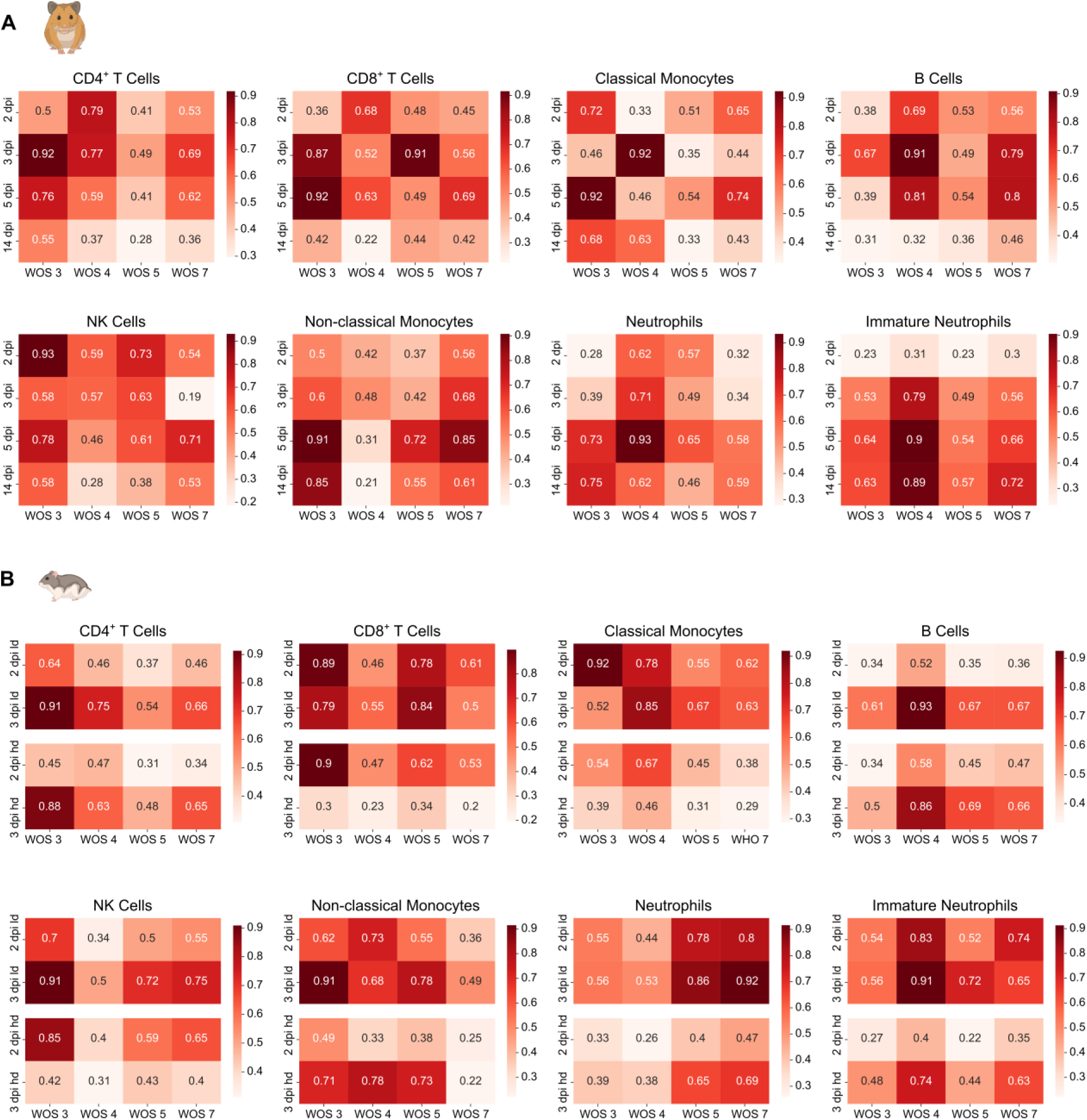
Neutrophils from Roborovski hamsters are the cell type that best matches that of patients with severe disease whereas Syrian hamster leukocytes are matched with moderate patient disease states. Overall cell type-specific transcriptome similarity from VAE pipeline for each time point in hamsters to each severity score of human patients quantified as similarity score (d2). Darker shades with higher values indicate higher similarity, corrected for species differences. (A) Similarity scores *d2* for pairs of human disease states (x-axis) and Syrian hamster disease states (y-axis). (B) Similarity scores d2 for pairs of human disease states (x-axis) and Roborovski hamster disease states (y-axis). WOS: WHO ordinal scale; dpi: days post infection; ld: low dose; hd: high dose.

### Differential gene expression correlation of human and hamster disease states

To examine which genes and pathways drive disease course similarities between COVID-19 patients and hamster models, we conducted a differential gene expression analysis of disease states compared to uninfected controls (**Supplementary Table4**). Globally, Syrian and Roborovski hamsters showed the highest percentage of differentially regulated genes 2dpi. In human patients, the highest percentage was observed in WOS7. To achieve the required minimum of two patients per group, WOS4 and WOS5 patients were analysed together. However, this group still displayed the lowest statistical power, as evident by the notably lower fraction of differentially regulated genes (Figure 5A, **Supplementary Table5**). We therefore focused on WOS3 and WOS7 and provide matching analyses for group WOS4 and WOS5 in the supplement (**Figure S5**). To identify similarities in antiviral responses, we next probed if significant differential regulation in human patients and hamsters occurs in the same direction (Figure 5B**, S5A**). On single gene level, effect-sizes of significant genes showed little correlation between humans and hamsters. Yet, classical monocytes showed the highest number of similarly regulated genes between WOS3 patients and 3dpi and 5dpi in Syrian hamsters, as well as with 2dpi in high-dose Roborovski hamsters. In contrast, for WOS7 patients, the monocyte gene expression resembled all acute phase time points (i.e. ≤5dpi) in both hamster species. Neutrophils showed the strongest concordance of gene effect sizes between WOS7 patients and 2dpi as well as 3dpi in Syrian and all Roborovski time points and doses, indicating increased activation of these cells in severe disease. In Syrian hamsters, neutrophils 5dpi showed more concordance with WOS3. At 14dpi, if at all, we only observed anti-correlated effect sizes of differentially regulated genes (e.g. CD4^+^-T-cells) between humans and Syrian hamsters. This was likely due to the resolution of the Syrian hamster’s infection by this point (Figure 5B). Notably, despite small correlation of differentially expressed genes, overlapping differentially expressed pathways between species were highly specific for COVID-19, relating to interferon signalling and inhibition of viral replication (Figure 5C**, S5B**). They showed the most prominent co-enrichment in neutrophils of WOS3 patients and Syrian as well as high-dose infected Roborovski hamsters. High co-enrichment was also observed in classical monocytes of WOS3 and WOS7 patients as well as Syrian and high-dose infected Roborovski hamsters (Figure 5C**, Supplementary Table6**).

**Figure 5.**
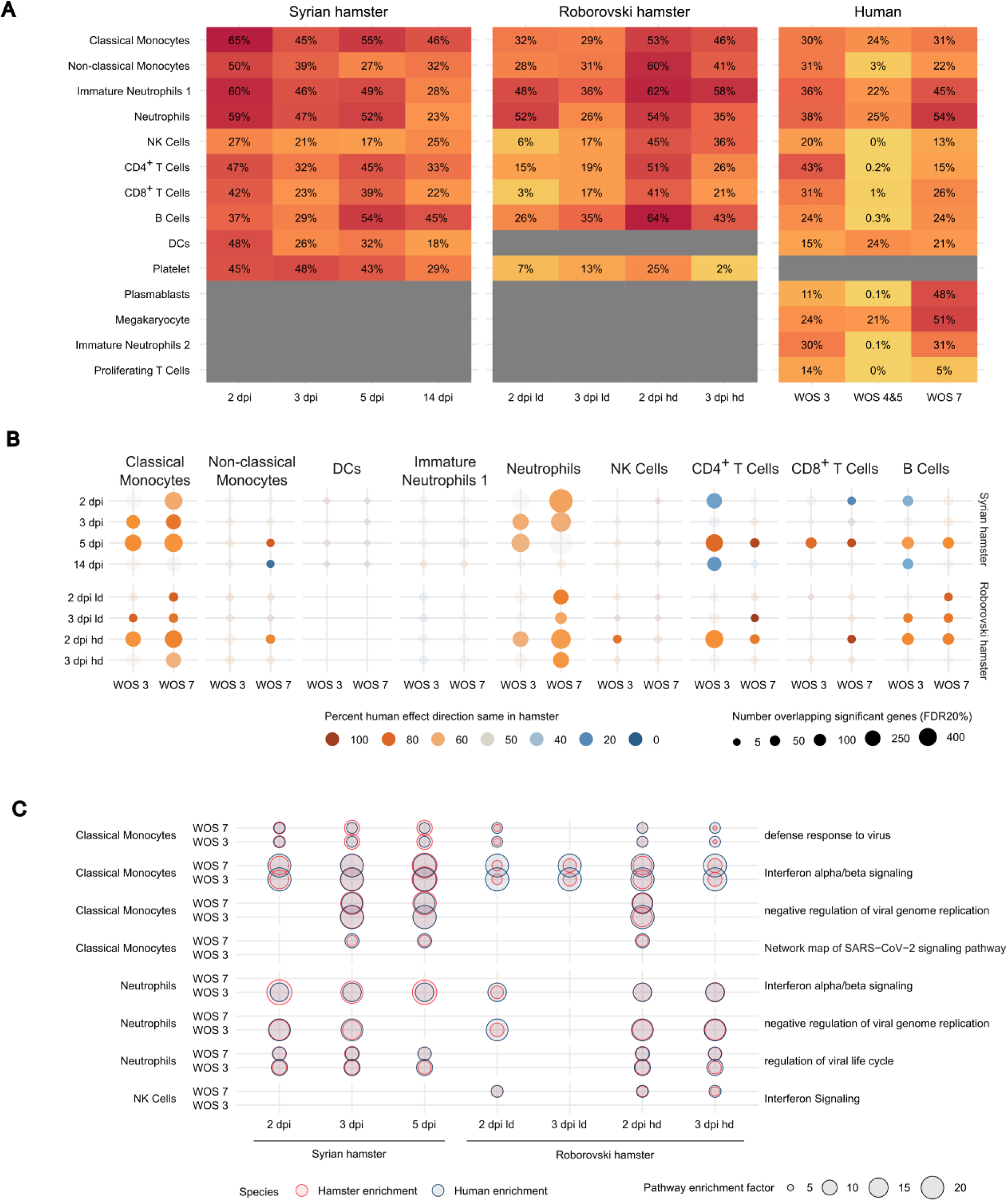
Differential gene expression correlation of human and hamster disease states. (A) Percentage of genes significantly up- or downregulated in humans or hamster species compared to controls. (B) Dotplot indicating significantly regulated genes regulated in the same direction in humans and hamsters. Shown in bright colors are conditions where significantly more (orange) or less (blue) than 50% of the overlapping genes were regulated in the same direction. (C) Dotplot of pathways enriched in differentially expressed genes that are present in human and hamster species. Here we focus on specific pathways, i.e. pathways not including more than 220 genes and not being redundant regarding included genes (see **Supplementary Table S5** for all pathways present in both human and hamsters). Differential expression analyses were performed in the Limma-Voom^35^ framework. Multiple testing adjustments were applied using the false-discovery-rate according to Benjamini and Hochberg. To correlate fold-changes, all genes significant at false discovery rate (FDR) ≤20% in both species were used. Pathway enrichment analyses for globally significant pathways (*P* corrected ≤ 0.05) were performed using gprofiler2. Pathway enrichment factor is the ratio between observed and expected proportion of significant genes in each pathway.

### Common most highly differentially expressed genes in humans and hamsters after infection

Based on the analysis of correlated genes, we sought to identify the top 10 genes for each cell type (ordered by absolute effect size at FDR 20%) that are regulated in matching directions in COVID-19 patients and hamsters (Figure 6**, S5C**). Interferon-regulated genes, like *IFI27*, *IFIT2,* and *IFIT3*, were most abundant during the early phase of infection in hamsters and matched both WOS3 and WOS7 patients across cell types. In neutrophils, all concordantly regulated top genes matched patients with severe disease progression. Most correlating neutrophil genes were expressed at 2dpi in Roborovski hamsters. *LCN2,* a regulator of interferon-stimulated gene expression, prominently correlated between high- and low-dose infected Roborovski hamsters and severe COVID-19 patients. Further, *HP* upregulation and *CSF1R* downregulation correlated in the early disease neutrophils of hamster models and severe patients. Interestingly, CD4^+^-T-cells and NK-cells displayed a pattern of most top regulated genes correlating between severe patients and Roborovski hamsters 2dpi of high-dose infection (Figure 6**, Supplementary Table7**).

**Figure 6.**
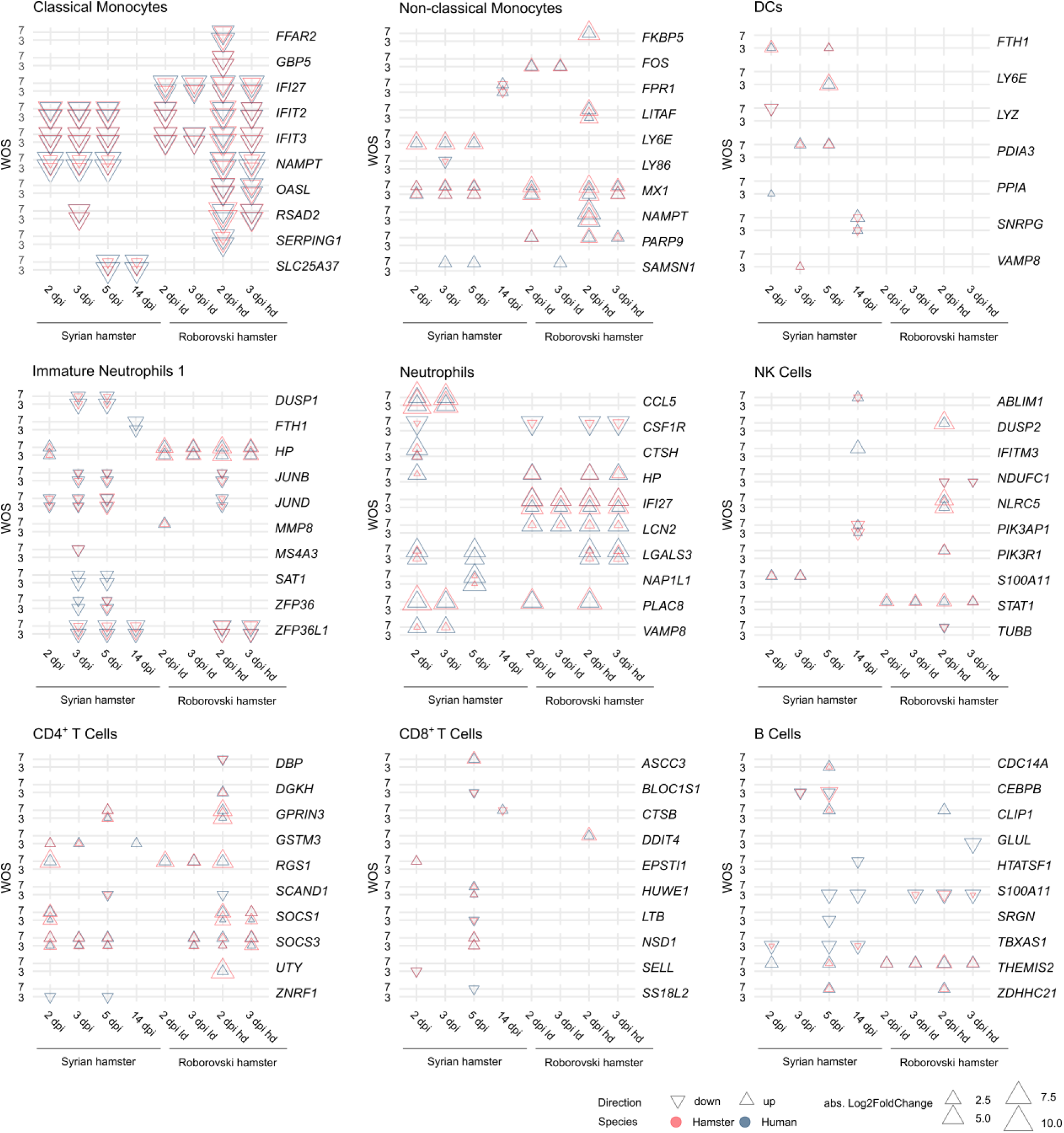
Common most differentially expressed genes in humans and hamsters after infection. Dotplot displaying the top 10 genes for each cell type (ordered by absolute effect size at FDR 20%) that are regulated in concordant direction (abs. Log2FoldChange) in humans and hamsters. Larger symbols correspond to larger effect sizes, direction of the triangles to up- or downregulation and color to species. All common significantly expressed genes with corresponding fold-changes are reported in **Supplementary Table S6.** Differential expression analyses were performed in the Limma-Voom^35^ framework. Multiple testing adjustments were applied using the false-discovery-rate (FDR) according to Benjamini and Hochberg.

### COVID-19 mediators and gene expression patterns amongst relevant cell types

To further investigate which specific genes contribute to severe or moderate disease courses, we analysed genes described to associated with inflammation^33,38^ (Figure 7A**, S6A, S6B, Supplementary Table8**). Several of these genes, such as interferon-stimulated genes (*IRF7*, *IFIT3*, *ISG15*, *ISG20*), chemokines (*CXCL10*, *CXCL11*) and pro-inflammatory mediators (*TNFSF10*, *NLRC5*) showed the highest expression levels in blood leukocytes of high-dose infected Roborovski hamsters. Low-dose infected Roborovski hamsters deteriorated less rapidly with fewer inflammatory genes expressed by classical monocytes. In human COVID-19 patients, the inflammatory response appeared more monocyte-dependent in moderate cases and neutrophil-dependent in severe cases. Interestingly, *S100A8*, *S100A9*, and *CD177* were not upregulated in hamsters, but were found to be upregulated in WOS7 patients and associate with severe COVID-19^10,39^ (Figure 7A).

**Figure 7.**
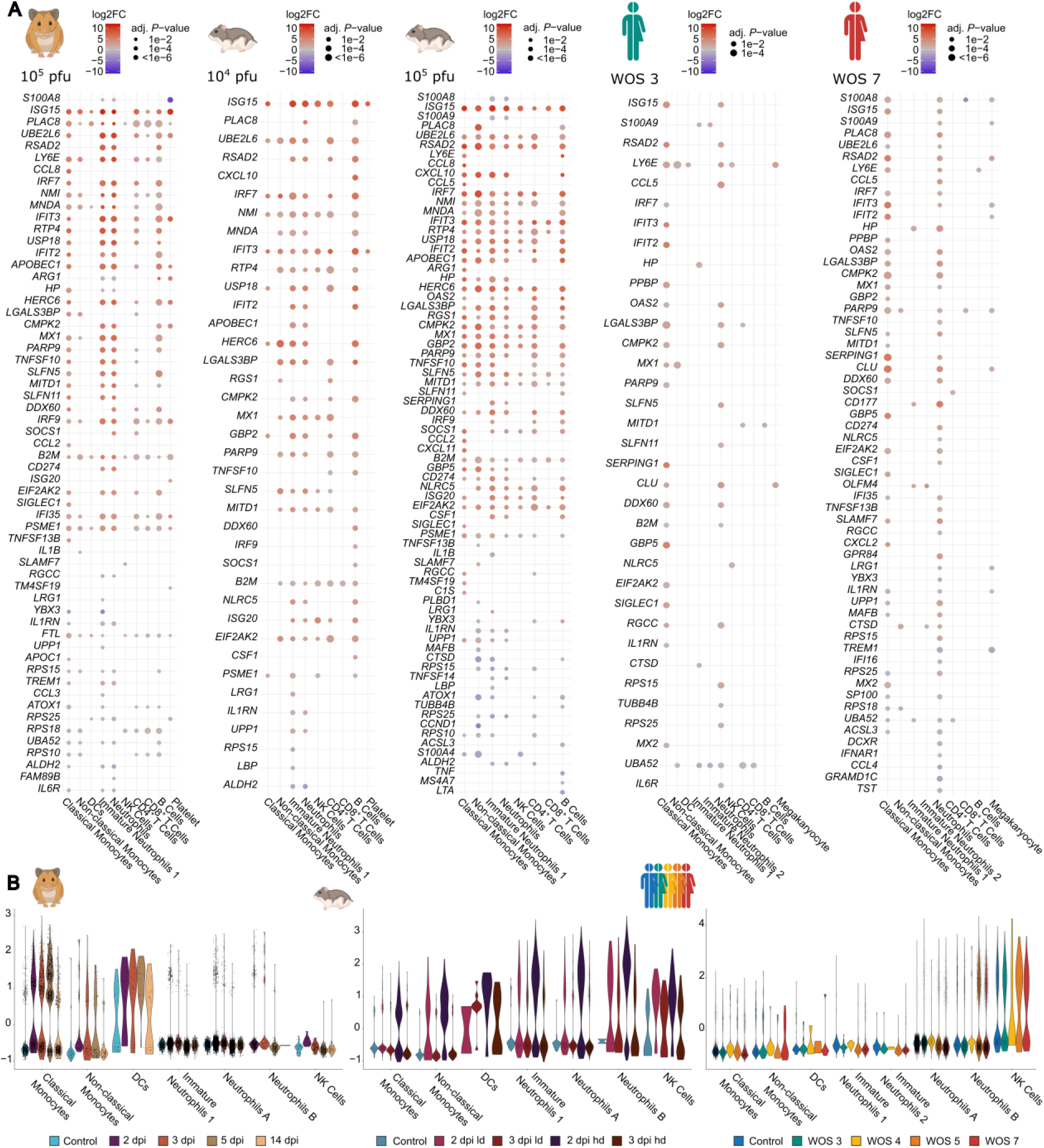
COVID-19 mediators and gene expression patterns amongst relevant cell types. (A) Dotplot displaying significantly up- or downregulated genes linked to inflammation in humans or hamster species. (B) Violin plots displaying expression of the previously described “severe inflammatory” gene set^40^ amongst different innate immune cell types in humans or hamster species. Differential expression analysis were performed in the Limma-Voom^35^ framework. Log2FC: log2FoldChange, adj. *P*-value: adjusted *P*-values were calculated according to Benjamini and Hochberg.

Finally, we analyzed a “severe inflammatory” gene signature in neutrophils linked to severe disease in humans^40^ (**Supplementary Table7**). This signature measures expression of 201 immune-related genes and was found in a subset of neutrophils of WOS5 and WOS7 COVID-19 patients. In Syrian hamsters, the gene scores in innate immune cells were low, whereas in Roborovski hamsters they were high, particularly in neutrophils of the high-dose infection group. Notably, this gene signature was not exclusively present in neutrophils, yet its severity-dependence was most prominent in neutrophils (Figure 7B).

While this demonstrates an inflammatory response of Roborovski hamster neutrophils similar to severe patients, a notable difference was that the gene set was found uniformly expressed in hamster neutrophils, but only in a specific subset of severe patients. (**Figure S7A**). Expression was high in the dense neutrophil-clusters “2” and “3” of severe patients, but low in the remaining neutrophil-clusters (**Figure S7A, S7B**). Differential expression analysis on clusters 2 and 3 compared to the other neutrophil clusters in severe patients further revealed elevated expression of inflammation-associated genes (**Figure S7C, S7D**). Notably, most of these genes were not included in the “severe inflammatory” signature and were yet highly expressed in “severe inflammatory” neutrophils (**Figure S7E**). In particular, genes associated with interferon response and TNFR signalling were highly expressed (**Figure S7C, S7D**). This suggests that the highly inflammatory response triggered by COVID-19 in human neutrophils is limited to a subset of these cells. Intriguingly, in Roborovski hamsters, the expression of the severe inflammatory signature was not restricted to a specific neutrophil subset.

## Discussion

Our study employs a novel computational approach^27^ to bridge the persistent translational divide between preclinical animal models and clinical research in humans^41^. By integrating VAE neural network-based generative models, differential gene expression, and pathway analysis, we achieved a thorough comparison of disease responses at a single-cell level across different species. Our VAE-based disease-state-matching pipeline demonstrates the feasibility of inferring humanized versions of hamster cells, addressing interspecies differences directly in the VAE latent space and extending existing cross-species integration strategies^42^.

Our VAE-based analysis corroborates that the Syrian hamster model closely resembles the response of human patients with moderate COVID-19 across all relevant blood cell types. Differential expression analysis confirmed this in specific, but highly relevant cell types: For example, the interferon-related gene expression peaking 5dpi in Syrian hamster classical monocytes resembles the immune response observed in immunocompetent patients^43^, while hamster neutrophils during early infection times match WOS4 patients with modest inflammatory response gene expression patterns. In sum, our approach further supports that SARS-CoV-2-infected Syrian hamsters are a valid model for studying innate and adaptive immunity in moderate COVID-19, in congruence with their proven effectiveness for testing vaccination^38^, immune modulatory therapy^44^, and antiviral therapeutic strategies^45,46^.

While Roborovski hamsters infected with high-dose SARS-CoV-2 have been utilized as a model for severe COVID-19 disease due to their high susceptibility^18,47^, our bioinformatics workflow shows some limitations: neutrophils are the only immune cell type in this model that reflects severe clinical disease in humans. Moreover, the rapid and fatal disease progression in Roborovski hamsters precludes the study of adaptive immunity during severe disease, as animals reach humane endpoints before cells like exhausted T-cells^6^ or pathologic CD16^+^ T-cells^48^ can develop. Besides, human patients with severe COVID-19 also exhibit increased inflammatory responses by innate immune cells other than neutrophils, which Roborovski hamsters fail to represent well, e.g. dysfunctional Eomes^hi^Tbet^lo^ NK-cells^49^. The same is true for the observed shift of inflammatory classical monocytes to “immunoparalysis” in severe COVID-19^6^. Multiple models may thus be required to simulate different aspects of severe human COVID-19, especially given the numerous risk factors influencing disease progression^50,51^. Hence, systematic approaches as proposed here might be a helpful tool also in other diseases to identify which processes in specific animal-cell types reflect the human states.

The neutrophil bias observed in Roborovski hamsters infected with SARS-CoV-2 is consistent across different infection doses, indicating a predisposition which was likewise observed in their pulmonary COVID-19 immune response^52^. This bias does not align with the diverse cellular inflammatory responses seen in humans with severe COVID-19 and some key genes associated with poor outcome in patients, namely the alarmins *S100A8*/*S100A9* encoding for calprotectin^10^ and the neutrophil activation marker *CD177^39^*, were not prominently expressed in hamster neutrophils. Consequently, the utility of the Roborovski hamster as a severe disease model for translating mechanisms of therapeutic immune modulators remains limited to subsets of inflammatory neutrophils^53^.

While offering valuable insights, our study has limitations. Sole reliance on blood data restricts organ-specific insights. Nonetheless, our focus on blood cells overcomes the problem that lung tissue is only available post-mortem from fatal cases, and oropharyngeal swabs as well as bronchoalveolar lavages, provide an incomplete picture of pulmonary cell types^54,55^. While our VAE neural network-based disease state matching offers unbiased scRNAseq data exploration, challenges arise in hyperparameter selection without a disease state ground truth in humans and hamsters. Careful handling of orthologues is crucial, as downstream analysis requires joint gene nomenclature between species. The VAE latent space lacks direct interpretability, although this can be mitigated by additional pathway and differential gene expression analyses to generate valuable biological insights. Furthermore, there is evidence that information from VAE has advantageous characteristics similar to PCA^56^. Additionally, a possible extension of our method is to consider linearly decoded VAE to provide a more direct link from latent variables of cells to genes^57^.

In summary, our innovative methodology supports the identification of meaningful animal models that represent human molecular dynamics for developing and testing therapeutic interventions. Our research helps to narrow the ‘translational gap’ by providing a nuanced understanding of disease responses across species at a single-cell level. It underscores the importance of critically evaluating any animal model to define its respective strengths and the limits of translatability, as demonstrated in the case of Roborovski hamsters as a supposed severe COVID-19 model.

## Acknowledgments

The authors thank Jeannine Wilde, Madlen Sohn, and Tatiana Borodina (MDC Scientific Genomics Platforms) for sequencing.

This work was supported by a German Federal Ministry of Education and Research (BMBF) grant to MS, MW, HK and GN, grant number e:Med CAPSyS (01ZX1304B, 01ZX1604B) and e:Med SYMPATH (01ZX1906A, 01ZX1906B); MW and GN were supported by the BMBF in the framework of MAPVAP (01KI2124). MW received funding from the DFG (project ID 114933180 – SFB-TR84, sub-projects C06 and C09 and project ID 431232613 – SFB-1449, sub-project B02). JT received funding from the Deutsche Forschungsgemeinschaft (DFG, German Research Foundation); project ID 114933180 – SFB-TR84, sub-project Z01b. VDF was financed by the Federal Ministry of Education and Research of Germany and by Sächsische Staatsministerium für Wissenschaft, Kultur und Tourismus in the programme Center of Excellence for AI-research „Center for Scalable Data Analytics and Artificial Intelligence Dresden/Leipzig“, project identification number: ScaDS.AI. AES thanks CompLS BMBF funding HOPARL. Schemes as well as Hamster and human icons were created using BioRender.com

## Authorship Contributions

Contribution: Conceptualization: VDF, PP, HK, GN; Methodology: VDF, PP, AES, EW, JT, HK, GN; Investigation: VDF, PP, EW, JMA, DP, LGTA, JK, FP, DV, TH, CG, ML, CG, AES, MS, MW, JT, HK, GN; Formal analysis: VDF, PP, HK, GN; Visualization: VDF, PP, HK, GN; Project administration: JT, HK, GN; Supervision: MS, MW, JT, HK, GN; Writing – original draft: VDF, PP, HK, GN

## References

1. Dinnon KH, 3rd, Leist SR, Okuda K, et al. SARS-CoV-2 infection produces chronic pulmonary epithelial and immune cell dysfunction with fibrosis in mice. Sci Transl Med. Sep 28 2022;14(664):eabo5070. doi:10.1126/scitranslmed.abo5070

2. Leenaars CHC, Kouwenaar C, Stafleu FR, et al. Animal to human translation: a systematic scoping review of reported concordance rates. J Transl Med. Jul 15 2019;17(1):223. doi:10.1186/s12967-019-1976-2

3. Vogel AB, Kanevsky I, Che Y, et al. BNT162b vaccines protect rhesus macaques from SARS-CoV-2. Nature. Apr 2021;592(7853):283-289. doi:10.1038/s41586-021-03275-y

4. Corbett KS, Flynn B, Foulds KE, et al. Evaluation of the mRNA-1273 Vaccine against SARS-CoV-2 in Nonhuman Primates. N Engl J Med. Oct 15 2020;383(16):1544–1555. doi:10.1056/NEJMoa2024671

5. Ferreira GS, Veening-Griffioen DH, Boon WPC, Moors EHM, van Meer PJK. Levelling the Translational Gap for Animal to Human Efficacy Data. Animals (Basel). Jul 15 2020;10(7)doi:10.3390/ani10071199

6. Su Y, Chen D, Yuan D, et al. Multi-Omics Resolves a Sharp Disease-State Shift between Mild and Moderate COVID-19. Cell. Dec 10 2020;183(6):1479–1495 e20. doi:10.1016/j.cell.2020.10.037

7. Schulte-Schrepping J, Reusch N, Paclik D, et al. Severe COVID-19 Is Marked by a Dysregulated Myeloid Cell Compartment. Cell. Sep 17 2020;182(6):1419–1440 e23. doi:10.1016/j.cell.2020.08.001

8. Sinha S, Rosin NL, Arora R, et al. Dexamethasone modulates immature neutrophils and interferon programming in severe COVID-19. Nat Med. Jan 2022;28(1):201–211. doi:10.1038/s41591-021-01576-3

9. Bernardes JP, Mishra N, Tran F, et al. Longitudinal Multi-omics Analyses Identify Responses of Megakaryocytes, Erythroid Cells, and Plasmablasts as Hallmarks of Severe COVID-19. Immunity. Dec 15 2020;53(6):1296–1314 e9. doi:10.1016/j.immuni.2020.11.017

10. Silvin A, Chapuis N, Dunsmore G, et al. Elevated Calprotectin and Abnormal Myeloid Cell Subsets Discriminate Severe from Mild COVID-19. Cell. Sep 17 2020;182(6):1401–1418 e18. doi:10.1016/j.cell.2020.08.002

11. Patel NG, Bhasin A, Feinglass JM, Angarone MP, Cohen ER, Barsuk JH. Mortality, critical illness, and mechanical ventilation among hospitalized patients with COVID-19 on therapeutic anticoagulants. Thrombosis Update. 2021/01/01/ 2021;2:100027. 10.1016/j.tru.2020.100027

12. Fan C, Wu Y, Rui X, et al. Animal models for COVID-19: advances, gaps and perspectives. Signal Transduct Target Ther. Jul 7 2022;7(1):220. doi:10.1038/s41392-022-01087-8

13. Ragan IK, Hartson LM, Dutt TS, et al. A Whole Virion Vaccine for COVID-19 Produced via a Novel Inactivation Method and Preliminary Demonstration of Efficacy in an Animal Challenge Model. Vaccines (Basel). Apr 1 2021;9(4)doi:10.3390/vaccines9040340

14. Lee JS, Koh JY, Yi K, et al. Single-cell transcriptome of bronchoalveolar lavage fluid reveals sequential change of macrophages during SARS-CoV-2 infection in ferrets. Nat Commun. Jul 27 2021;12(1):4567. doi:10.1038/s41467-021-24807-0

15. Speranza E, Williamson BN, Feldmann F, et al. Single-cell RNA sequencing reveals SARS-CoV-2 infection dynamics in lungs of African green monkeys. Sci Transl Med. Jan 27 2021;13(578)doi:10.1126/scitranslmed.abe8146

16. Qin Z, Liu F, Blair R, et al. Endothelial cell infection and dysfunction, immune activation in severe COVID-19. Theranostics. 2021;11(16):8076–8091. doi:10.7150/thno.61810

17. Imai M, Iwatsuki-Horimoto K, Hatta M, et al. Syrian hamsters as a small animal model for SARS-CoV-2 infection and countermeasure development. Proc Natl Acad Sci U S A. Jul 14 2020;117(28):16587–16595. doi:10.1073/pnas.2009799117

18. Trimpert J, Vladimirova D, Dietert K, et al. The Roborovski Dwarf Hamster Is A Highly Susceptible Model for a Rapid and Fatal Course of SARS-CoV-2 Infection. Cell Rep. Dec 8 2020;33(10):108488. doi:10.1016/j.celrep.2020.108488

19. Gruber AD, Firsching TC, Trimpert J, Dietert K. Hamster models of COVID-19 pneumonia reviewed: How human can they be? Vet Pathol. Jul 2022;59(4):528–545. doi:10.1177/03009858211057197

20. Luecken MD, Buttner M, Chaichoompu K, et al. Benchmarking atlas-level data integration in single-cell genomics. Nat Methods. Jan 2022;19(1):41–50. doi:10.1038/s41592-021-01336-8

21. Sikkema L, Ramirez-Suastegui C, Strobl DC, et al. An integrated cell atlas of the lung in health and disease. Nat Med. Jun 2023;29(6):1563–1577. doi:10.1038/s41591-023-02327-2

22. Xu C, Lopez R, Mehlman E, Regier J, Jordan MI, Yosef N. Probabilistic harmonization and annotation of single-cell transcriptomics data with deep generative models. Mol Syst Biol. Jan 2021;17(1):e9620. doi:10.15252/msb.20209620

23. Kingma DP, Welling M. Auto-Encoding Variational Bayes. presented at: ICLR; 2014; http://dblp.uni-trier.de/db/conf/iclr/iclr2014.html#KingmaW13kingma2014autoencoding

24. Kingma DP, Welling M. An Introduction to Variational Autoencoders. Foundations and Trends® in Machine Learning. 2019;12(4):307–392. doi:10.1561/2200000056

25. Gronbech CH, Vording MF, Timshel PN, Sonderby CK, Pers TH, Winther O. scVAE: variational auto-encoders for single-cell gene expression data. Bioinformatics. Aug 15 2020;36(16):4415–4422. doi:10.1093/bioinformatics/btaa293

26. De Donno C, Hediyeh-Zadeh S, Moinfar AA, et al. Population-level integration of single-cell datasets enables multi-scale analysis across samples. Nat Methods. Nov 2023;20(11):1683–1692. doi:10.1038/s41592-023-02035-2

27. Lotfollahi M, Wolf FA, Theis FJ. scGen predicts single-cell perturbation responses. Nat Methods. Aug 2019;16(8):715–721. doi:10.1038/s41592-019-0494-8

28. Heydari AA, Davalos OA, Zhao L, Hoyer KK, Sindi SS. ACTIVA: realistic single-cell RNA-seq generation with automatic cell-type identification using introspective variational autoencoders. Bioinformatics. Apr 12 2022;38(8):2194–2201. doi:10.1093/bioinformatics/btac095

29. Lotfollahi M, Naghipourfar M, Theis FJ, Wolf FA. Conditional out-of-distribution generation for unpaired data using transfer VAE. Bioinformatics. Dec 30 2020;36(Suppl_2):i610-i617. doi:10.1093/bioinformatics/btaa800

30. Lotfollahi M, Naghipourfar M, Luecken MD, et al. Mapping single-cell data to reference atlases by transfer learning. Nat Biotechnol. Jan 2022;40(1):121–130. doi:10.1038/s41587-021-01001-7

31. Tasaki S, Xu J, Avey DR, et al. Inferring protein expression changes from mRNA in Alzheimer’s dementia using deep neural networks. Nat Commun. Feb 3 2022;13(1):655. doi:10.1038/s41467-022-28280-1

32. Kana O, Nault R, Filipovic D, Marri D, Zacharewski T, Bhattacharya S. Generative modeling of single-cell gene expression for dose-dependent chemical perturbations. Patterns (N Y). Aug 11 2023;4(8):100817. doi:10.1016/j.patter.2023.100817

33. Nouailles G, Wyler E, Pennitz P, et al. Temporal omics analysis in Syrian hamsters unravel cellular effector responses to moderate COVID-19. Nat Commun. Aug 11 2021;12(1):4869. doi:10.1038/s41467-021-25030-7

34. Haghverdi L, Buttner M, Wolf FA, Buettner F, Theis FJ. Diffusion pseudotime robustly reconstructs lineage branching. Nat Methods. Oct 2016;13(10):845–8. doi:10.1038/nmeth.3971

35. Law CW, Chen Y, Shi W, Smyth GK. voom: Precision weights unlock linear model analysis tools for RNA-seq read counts. Genome Biol. Feb 3 2014;15(2):R29. doi:10.1186/gb-2014-15-2-r29

36. Pennitz P, Kirsten H, Friedrich VD, et al. A pulmonologist’s guide to perform and analyse cross-species single lung cell transcriptomics. Eur Respir Rev. Sep 30 2022;31(165)doi:10.1183/16000617.0056-2022

37. Hao Y, Stuart T, Kowalski MH, et al. Dictionary learning for integrative, multimodal and scalable single-cell analysis. Nat Biotechnol. May 25 2023;doi:10.1038/s41587-023-01767-y

38. Nouailles G, Adler JM, Pennitz P, et al. Live-attenuated vaccine sCPD9 elicits superior mucosal and systemic immunity to SARS-CoV-2 variants in hamsters. Nat Microbiol. May 2023;8(5):860–874. doi:10.1038/s41564-023-01352-8

39. Levy Y, Wiedemann A, Hejblum BP, et al. CD177, a specific marker of neutrophil activation, is associated with coronavirus disease 2019 severity and death. iScience. Jul 23 2021;24(7):102711. doi:10.1016/j.isci.2021.102711

40. Aschenbrenner AC, Mouktaroudi M, Kramer B, et al. Disease severity-specific neutrophil signatures in blood transcriptomes stratify COVID-19 patients. Genome Med. Jan 13 2021;13(1):7. doi:10.1186/s13073-020-00823-5

41. Seyhan AA. Lost in translation: the valley of death across preclinical and clinical divide – identification of problems and overcoming obstacles. Translational Medicine Communications. 2019/11/18 2019;4(1):18. doi:10.1186/s41231-019-0050-7

42. Song Y, Miao Z, Brazma A, Papatheodorou I. Benchmarking strategies for cross-species integration of single-cell RNA sequencing data. Nat Commun. Oct 14 2023;14(1):6495. doi:10.1038/s41467-023-41855-w

43. Maher AK, Burnham KL, Jones EM, et al. Transcriptional reprogramming from innate immune functions to a pro-thrombotic signature by monocytes in COVID-19. Nat Commun. Dec 26 2022;13(1):7947. doi:10.1038/s41467-022-35638-y

44. Rocha SM, Fagre AC, Latham AS, et al. A Novel Glucocorticoid and Androgen Receptor Modulator Reduces Viral Entry and Innate Immune Inflammatory Responses in the Syrian Hamster Model of SARS-CoV-2 Infection. Front Immunol. 2022;13:811430. doi:10.3389/fimmu.2022.811430

45. Abdelnabi R, Foo CS, Jochmans D, et al. The oral protease inhibitor (PF- 07321332) protects Syrian hamsters against infection with SARS-CoV-2 variants of concern. Nat Commun. Feb 15 2022;13(1):719. doi:10.1038/s41467-022-28354-0

46. Maio N, Cherry S, Schultz DC, Hurst BL, Linehan WM, Rouault TA. TEMPOL inhibits SARS-CoV-2 replication and development of lung disease in the Syrian hamster model. iScience. Oct 21 2022;25(10):105074. doi:10.1016/j.isci.2022.105074

47. Lieber CM, Cox RM, Sourimant J, et al. SARS-CoV-2 VOC type and biological sex affect molnupiravir efficacy in severe COVID-19 dwarf hamster model. Nat Commun. Jul 29 2022;13(1):4416. doi:10.1038/s41467-022-32045-1

48. Georg P, Astaburuaga-Garcia R, Bonaguro L, et al. Complement activation induces excessive T cell cytotoxicity in severe COVID-19. Cell. Feb 3 2022;185(3):493–512 e25. doi:10.1016/j.cell.2021.12.040

49. Witkowski M, Tizian C, Ferreira-Gomes M, et al. Untimely TGFbeta responses in COVID-19 limit antiviral functions of NK cells. Nature. Dec 2021;600(7888):295-301. doi:10.1038/s41586-021-04142-6

50. Richardson S, Hirsch JS, Narasimhan M, et al. Presenting Characteristics, Comorbidities, and Outcomes Among 5700 Patients Hospitalized With COVID-19 in the New York City Area. JAMA. May 26 2020;323(20):2052-2059. doi:10.1001/jama.2020.6775

51. Benitez ID, de Batlle J, Torres G, et al. Prognostic implications of comorbidity patterns in critically ill COVID-19 patients: A multicenter, observational study. Lancet Reg Health Eur. Jul 2022;18:100422. doi:10.1016/j.lanepe.2022.100422

52. Peidli S, Nouailles G, Wyler E, et al. Single-cell-resolved interspecies comparison identifies a shared inflammatory axis and a dominant neutrophil-endothelial program in severe COVID-19. bioRxiv. 2023:2023.08.25.551434. doi:10.1101/2023.08.25.551434

53. Wyler E, Adler JM, Eschke K, et al. Key benefits of dexamethasone and antibody treatment in COVID-19 hamster models revealed by single-cell transcriptomics. Mol Ther. May 4 2022;30(5):1952–1965. doi:10.1016/j.ymthe.2022.03.014

54. Chua RL, Lukassen S, Trump S, et al. COVID-19 severity correlates with airway epithelium-immune cell interactions identified by single-cell analysis. Nat Biotechnol. Aug 2020;38(8):970–979. doi:10.1038/s41587-020-0602-4

55. Grant RA, Morales-Nebreda L, Markov NS, et al. Circuits between infected macrophages and T cells in SARS-CoV-2 pneumonia. Nature. Feb 2021;590(7847):635-641. doi:10.1038/s41586-020-03148-w

56. Rolinek M, Zietlow D, Martius G. Variational Autoencoders Pursue PCA Directions (by Accident). 2019 IEEE/CVF Conference on Computer Vision and Pattern Recognition (CVPR). 2018:12398–12407.

57. Virshup I, Bredikhin D, Heumos L, et al. The scverse project provides a computational ecosystem for single-cell omics data analysis. Nat Biotechnol. May 2023;41(5):604–606. doi:10.1038/s41587-023-01733-8

58. Lun ATL, Richard AC, Marioni JC. Testing for differential abundance in mass cytometry data. Nat Methods. Jul 2017;14(7):707–709. doi:10.1038/nmeth.4295

