## Supplemental Material and Supplemental Figures for "Neural Network-Assisted Humanization of COVID-19 Hamster scRNAseq Data Reveals Matching Severity States in Human Disease"

### Supplemental Methods

#### Virus stocks

SARS-CoV-2 isolate (BetaCoV/Germany/BavPat1/2020)[1] was kindly provided by Daniela Niemeyer and Christian Drosten, Charité Berlin, Germany. The virus stocks required for animal experiments were generated by propagating the virus under BSL-3 conditions on Vero E6 cells (ATCC CRL-1586) in minimal essential medium (MEM; PAN Biotech, Aidenbach, Germany) supplemented with 10% fetal bovine serum (FBS , PAN), 100 IU/ml penicillin G and 100 µg/ml streptomycin (Carl Roth, Karlsruhe, Germany). Low-passage virus stocks used for animal experiments were titrated on Vero-E6 cells before the start of infection experiments. Briefly, Vero E6 cells were incubated with serial 10-fold dilutions of virus stocks for 2 h. The virus inoculum was replaced with an overlay medium consisting of Dulbecco's modified Eagle's medium (DMEM, PAN Biotech, Aidenbach, Germany), 2.5% microcrystalline cellulose (Avicel RC-591, DuPont, Wilmington, DE, USA) and 10% fetal bovine serum (FBS PAN Biotech, Aidenbach, Germany). Following 72 hours of incubation at 37°C in a 5% CO2 atmosphere, inoculated cells were fixed with 4% formaldehyde for 24 hours. Virus induced cytopathic effect (plaques) were visualized by methylene blue counterstaining.

The genome sequence integrity of virus stocks used for animals experiments was determined by Illumina sequencing as described[2] and aligned to the reference sequence of the isolate (GenBank: MT270101 and GISAID: EPI_ISL_406862). Specifically, the presence and integrity of the furin cleavage site within the viral spike glycoprotein was confirmed by sequencing.

**Animals and Infection**

In application of the 3R principle, no separate animals were sacrificed for the purpose of this study. Instead, we re-used data (Syrian hamsters[3]) and blood samples (Roborovski hamsters[4,5]) from animals studied in our previously published research. As part of these experiments, female and male Syrian hamsters (*Mesocricetus auratus*; RjHan:AURA, Janvier Labs, Saint-Berthevin, France) and Roborovski hamsters (*Phodopus roborovskii*, German pet store) were kept in individually ventilated cages (Tecniplast, Buguggiate, Italy) in a BSL-3 laboratory with environmental enrichment (Carfil, Oud-Turnhout, Belgium) and free access to food and water. For the duration of the infection experiments, the cage temperature and relative humidity were set between 22 and 24 °C and 40 and 55%, respectively. Before each experiment, the animals were allowed to acclimatize to the experimental conditions for at least 7 days. As described in previous work[6], 10- to 12-week-old male and female Syrian and Roborovski hamsters were anesthetized and intranasally infected with $1 \times 10^5$ or $1 \times 10^4$ plaque-forming units (pfu) SARS-CoV-2 (variant B1, isolate BetaCoV /Germany)/BavPat1/ 2020). We performed clinical examination of all infected animals twice daily. Animals with a body weight loss of more than 15% for more than 48 h were euthanized according to the animal use protocol. In all other instances, naive hamsters (n = 3) and hamsters at 2, 3, 5 and 14 days after infection (n = 3 each) were randomly selected for sample collection. As described in our previous work[4], euthanasia was carried out by cervical dislocation and exsanguination under anaesthesia. For the purpose of this study, the collection of approx. 1 ml of EDTA anti-coagulated blood for transcriptome and single-cell transcriptome studies was particularly relevant[3]. The hamster blood sampling schedule reflected their disease course: Roborovski hamsters experience a severe to fatal disease course, meeting humane endpoint criteria at 3 days post-infection (dpi), while Syrian hamsters undergo a transient, moderate disease course that allows for follow-up analyses of virus clearance and resolution of inflammation after high-dose infection. Thus, additional 5 dpi and 14 dpi time-points were sampled for Syrian hamsters. For human data, actual time-point of infection was unknown. Human data were grouped by WOS quantifying disease severity, and included patients with WOS3, WOS4, WOS5 and WOS7.

**Isolation of white blood cells and single-cell transcriptome library generation**

500 µL whole blood per animal was lysed in red blood cell lysis buffer (ChemCruz), washed and centrifuged according to the manufacturer’s instructions. Each resulting RBC-free pellet was resuspended in low-BSA buffer (1× PBS, 0.04% BSA), filtered with 40 µm FloMi filters (Merck) and the number of cells determined using a hemocytometer and trypan blue to estimate the fraction of apoptotic cells. Isolated cells were then processed for single-cell RNA sequencing according to the manufacturer’s instructions (Chromium Next GEM Single Cell 3' Reagent Kit v3.1; 10x Genomics, Pleasanton, CA, USA). The resulting single-cell libraries were quantified using Qubit (ThermoFisher) and quality-controlled using the Bioanalyzer system (Agilent). Sequencing was performed on a Novaseq 6000 device (Illumina) according to the manufacturer’s instructions.

**Species-specific processing of data**

*Phodopus roborovskii* raw sequencing files were processed using Cellranger mkfastq (10x Genomics). Fastq files were aligned to the Roborovski hamster genome assembly PHOROB[7], filtered and counted using cell ranger count (10x Genomics). R (Version 4.2.2) was used to analyze the raw feature-barcode matrix, including the R-package SeuratV5 4.9.9.9044 for single-cell analysis[8].

*Mesocricetus auratus* expression sequencing data aligned to a modified version of Ensembl 99 MesAur 1.0 were downloaded as Seurat-objects[3]. The human phenotype and single-cell data were downloaded as Seurat-objects[9] and gene counts were corrected for technical batches using method Combat. Disease severity of COVID-19 patients was estimated using WOS. WOS1 and WOS2 were considered mild, WOS3 and WOS4 moderate, and WOS5 to WOS7 severe[10]. On each dataset of the three species, the following quality filtering steps were performed separately: First, ambient RNA was removed using DecontX framework from R package celda 1.14.2 and DropletUtils 1.16.0, and doublets were identified by combining predictions from DoubletFinder 2.0.3 and scDblFinder1.13.12. The 4,000 most variable genes were used in principal component analysis to reduce dimensionality and define clusters of cells at a resolution of 0.8. Next, we identified clusters where more than 50% of cells had atypical numbers of gene counts, quantified as unique molecular identifier (UMIs), of different genes expressed, of mitochondrial, ribosomal, and hemoglobin-related genes, where the most abundant 50 different UMIs accounted for a considerable proportion of all counts or where cells were marked as doublets (See **Supplementary Table 1** for cut-off values). Finally, using a more stringent cutoff, we identified individual cells not meeting our quality control criteria (See **Supplementary Table 1**) or being marked as doublets. This defined a total of 6.2%, 8.7%, and 4.1% of cells from

*Phodopus roborovskii*, *Mesocricetus auratus,* and human data, as low quality cells, respectively.

**VAE model**

We applied a variational autoencoder (VAE) model[11] for jointly embedding high-dimensional hamster and human gene expression data into a lower dimensional latent space. The VAE model consists of two main components - an encoder neural network and a decoder neural network. The encoder takes as an input the original gene expression data and maps it to a lower-dimensional latent space. The mapping is achieved by forward propagation of the input through the hidden layers of the encoder. The final layer of the encoder is referred to as the ’bottleneck layer’ and consists of fewer nodes compared to the input. The decoder takes the low-dimensional output of the encoder as an input and tries to reconstruct the original input data as precisely as possible, a process referred to as decoding. By forcing the high-dimensional input data to pass through a low-dimensional bottleneck, only the most relevant and meaningful structures are captured in latent space. A VAE not only provides a latent embedding but also learns a multivariate distribution in latent space that best describes the embedded data and allows for data generation[12]. Implementation was carried out in a python v3.9.5 and scanpy[13] v1.8.2 environment. The python tool scGen[11] v2.0.0 was used for VAE implementation and execution (*scgen.SCGEN*). The scGen VAE architecture is especially designed for the analysis and prediction of gene expression across conditions and species[11]. Separate VAE models were trained for the comparison between humans and Syrian hamsters as well as between humans and Roborovski hamsters. Furthermore, to account for diverse transcriptomic responses to infection across cell types, distinct VAE models were learned at the cell type level. Cell types that contained enough cells in various severity groups of human patients or time points in hamster data to train a VAE were investigated: Classical monocytes, non-classical monocytes, neutrophils, immature neutrophils, $CD4^+$-T-cells, $CD8^+$-T-cells, B-cells and NK-cells. The gene counts of combined highly variable hamster genes and highly variable human genes were used as input to the VAE. These harbor the most relevant biological signals while avoiding excessive input burden on the VAE. Highly variable genes were computed with *scanpy.pp.highly_variable_genes* and default parameters. The encoder and decoder networks of the VAE both consisted of two hidden layers with 800 neurons. The latent space dimension was set to 10 with an underlying Gaussian latent distribution. An overview of the used scGen model parameters can be found in section **VAE model parameters**.

#### VAE training

Training of the VAE model was carried out with scGen method *scgen.SCGEN.train.* Training parameters are shown in section **VAE model parameters***.* To account for stochasticity in the training procedure, five VAE models were trained per hamster species and cell type for disease state matching. The model that promised the highest separability of disease states in 10-dimensional latent space was finally selected for disease state matching. To this end, we fitted a MANOVA model[14] making use of the python package statsmodels[15] v0.13.1 with disease state label as dependent variable and the 10 latent dimensions as independent variables. The Pillai's trace of the MANOVA model served as heuristic for disease state separability in latent space. For each cell type and hamster species, we selected the model with the highest Pillai's trace for disease state matching.

**VAE latent space arithmetic**

Hamster and human data was mapped from high-dimensional gene expression space into 10-dimensional latent space via the trained encoder of the VAE model. To address general interspecies difference in latent space, the species-shift-vector $\delta_{species}$ was computed based on human controls and hamster controls. The exclusive use of controls in this step ensured that the species-shift-vector $\delta_{species}$ controlled for interspecies difference while not being affected by infection response. Technically, the species-shift-vector $\delta_{species}$ was obtained by subtracting the mean latent embedding of hamster controls from the mean latent embedding of human controls. To allow for cross-species disease state comparison, the species-shift-vector $\delta_{species}$ was then applied to the latent embeddings of infectious hamster cells in VAE latent space which we refer to as humanization.

**VAE similarity measure**

The distance between humanized hamster disease states and human disease states in 10-dimensional latent space was quantified via diffusion pseudotime[16] (dpt). Dpt is a graph-based distance metric that offers a (pseudo)temporal ordering of cells relative to a root cell. We made use of dpt as distance measure in the following manner: For each humanized hamster disease state, a representative root cell was selected at random. Dpt distance between the latent embedding of the humanized hamster root cell and the latent embeddings of infectious human cells was computed. The mean dpt distance within a human disease state to the humanized hamster root cell was used as proxy for the similarity of that human disease state to the humanized hamster root cell. This procedure was repeated and averaged over 10 different humanized hamster root cells. Low dpt distance indicates high similarity while high dpt distance indicates low similarity. For better intuition, we scaled and transformed the mean dpt outcomes to the range from 0 to 1 where values close to 1 indicate high similarity and values close to 0 signal low similarity. Details can be found in **Mathematical notation – similarity score**. As the graph-based dpt distance is dependent on the root cell, we repeated the described procedure with human root cells. The final similarity between a human disease state and a humanized hamster disease state is then based on dpt distances with human and humanized hamster root cells, both equally weighted. Dpt distance was calculated via *scanpy.tl.dpt* with default parameters.

**Cross-species proof of principle**

For the cross-species proof of principle, we trained one VAE model for human and Syrian hamster neutrophil controls as well as one VAE model for human and Roborovski hamster non-classical monocyte controls (**Figure 3 A, B**). We performed a training-test split the following way: The training set included two out of three hamster replicates and all but one of the human donors. The test set comprised the excluded hamster replicate (*Ha1*) and human donor (*BN-31*). The species-shift-vector $\delta_{species}$ was derived de novo solely from the VAE latent embeddings of the training set as described in **VAE latent space arithmetic**. The hamster test set was then embedded using the VAE encoder from the training set. Furthermore, we added $\delta_{species}$ from the training set to each hamster test set latent cell representation to humanize the embedded hamster controls. Following this, we mapped the latent representations back to gene expression space via the VAE decoder from the training set.

**Intra-species proof of principle**

For the intra-species proof of principle, we trained one VAE model for Syrian hamster neutrophils as well as one VAE model for Roborovski hamster non-classical monocytes (**Figure 3C, 3D**). We performed a training-test split the following way: The training set included two out of three hamster replicates while the test set consisted of the remaining hamster replicate *Ha1*. The infection-shift-vector $\delta_{infection}$ was derived on the training set by subtracting the mean latent embedding of the respective control groups from the mean latent embedding of 2dpi cells (Syrian hamster) or high-dose 2dpi cells (Roborovski hamster). The test set was then embedded using the VAE encoder from the training set. $\delta_{infection}$ was added to the latent representations of the test control cells to predict a diseased version of the respective control cells in VAE latent space. Eventually, the VAE decoder from the training set was used for mapping the latent representation back to gene expression space.

#### VAE model parameters

The following parameters were used for implementation of the VAE model with *scgen.SCGEN* class: *n_hidden*: 800; *n_latent*: 10; *dropout_rate*: 0.2. For training of the VAE model the *scgen.SCGEN.train* method was applied with following parameters: *max_epochs*: 100; *use_gpu*: None; *train_size*: 0.9; *batch_size*: 32; *early_stopping*: True; *early_stopping_patience*: 25.

#### Tools in VAE implementation

We used the following main tools within the VAE disease state matching pipeline: Python (version 3.9.5); scGen[11] (version 2.0.0); scVI[17] (version 0.13.0); Scanpy[13] (version 1.8.2); NumPy[18] (version 1.21.5); Seaborn[19] (version 0.11.2); pandas[20] (version 1.4.0); AnnData[21] (version 0.9.1); PyTorch[22] (version 1.10.2+cu102); SciPy[23] (version 1.8.0); Matplotlib[24] (version 3.5.1); statsmodels[15] (version 0.13.1).

**Mathematical notation - similarity score**

Let *D* denote the data under consideration. We define with

$$dpt(r,c)$$

the diffusion pseudotime distance between root cell $r$ and any other cell $c$. Let $P \subset D$ denote a subpopulation of $D$. We define the mean diffusion pseudotime distance between root *r* and $P$ as

$$\overline{dpt}(r,P) \ := \frac{1}{|P|}\sum_{p\epsilon P} dpt(r,p)$$

for $|P| > 0$. Let *B* denote a base population. We sample $K$ different root cells $r_1,...,r_K$ from $B$. We define

$$dptm(B,P) \ := \frac{1}{K}\sum_{j=1}^{K} \overline{dpt}(r_j,P)$$

as the mean diffusion pseudotime distance between $B$ and $P$ with root cells taken from $B$. Assume we compute $dptm$ for a base population $B$ and $C > 0$ subpopulations $P_1,...,P_C$. We define

$$max \coloneqq \max_{c\epsilon\,\{1,\ldots,C\}} dptm(B,P_c)$$

$$min \coloneqq \min_{c\epsilon\,\{1,\ldots,C\}} dptm(B,P_c)$$

$$L(B,P_d) := dptm(B,P_d) - min + 0.1(max - min)$$

$$Sim(B,P_d) \ := 1 - \frac{L(B,P_d)}{\sum_{c\epsilon C} L(B,P_c)}$$

Let $Hum_1,...,Hum_q$ denote $Q > 0$ human subpopulations and $Ha_1,...,Ha_J$ denote $J > 0$ hamster subpopulations. $Sim$ is a directed similarity measure as graph root cells are drawn from the base population $B$. To obtain an undirected similarity measure $d_2$ for a human disease state $Hum_q$ and a hamster disease state $Ha_j$, we consider root cells from both human and hamster subpopulations in the following manner:

$$d_2(Hum_q , Ha_j) := Sim(Hum_q , Ha_j) \times Sim(Ha_j, Hum_q)$$

By construction we have that

$$d_2(Hum_q,Ha_j) = d_2(Ha_j,Hum_q).$$

We make use of $d_2$ as distance metric between two disease states.

### Mathematical notation - variational autoencoder (VAE)

We denote the encoder part of the VAE model as $enc$ and the decoder part as $dec$. For $M$−dimensional gene expression data and $L$−dimensional latent space, the encoder can be compactly summarized as a non-linear mapping

$$enc : R^M \rightarrow R^L$$

whereas the decoder can be summarized as a non-linear mapping of the form

$$dec : R^L \rightarrow R^M.$$

### Mathematical notation - latent space arithmetics

Assume we have trained the VAE model on $Hum = Hum^{control} \cup Hum^{infection}$ and $Ha = Ha^{control} \cup Ha^{infection}$ human and hamster $M$−dimensional gene expression data for a specific cell type and hamster species. Let

$$Z_{human}{}^{control} := \{enc(x) \in \mathrm{R}^L | x \in Hum^{control}\}$$

$$Z_{hamster}{}^{control} := \{enc(x) \in \mathrm{R}^L | x \in Ha^{control}\}$$

denote the latent embeddings of the human and hamster control groups. We compute the species-shift-vector $\delta_{species}$ as

$$\delta_{species} := mean(Z_{human}^{control}) - mean(Z_{hamster}^{control}).$$

Note that $\delta_{species} \in \mathrm{R}^{L}$ is solely based on human and hamster controls. Let

$$Z_{hamster}^{infection} := \{enc(x) \in R^{L} | x \in Ha^{infection}\}$$

denote the latent embedding of hamster infection group. We humanize $Z_{hamster}^{infection}$ by applying $\delta_{species}$ the following way:

$$\overline{Z}_{hamster}^{infection} := Z_{hamster}^{infection} + \delta_{species} = \{enc(x) + \delta_{species} | x \in Ha^{infection}\}.$$

Let further

$$Z_{human}^{infection} := \{enc(x) \in R^{L} | x \in Hum^{infection}\}$$

denote the latent embedding of the human infection group. Disease state matching is then based on the humanized latent hamster embedding $\overline{Z}_{hamster}^{infection}$ and the latent human embedding $Z_{human}^{infection}$. Assume we have four different humanized hamster embedded subpopulations $\overline{Ha}_{1},\ldots,\overline{Ha}_{4}$ with $\overline{Ha}_{j} \subset \overline{Z}_{hamster}^{infection}$ for $1 \leq j \leq 4$ and four different human embedded subpopulations $Hum_{1},\ldots,Hum_{4}$ with $Hum_{q} \subset Z_{human}^{infection}$ for $1 \leq q \leq 4$. Similarity between $\overline{Ha}_{j}$ and $Hum_{q}$ is then evaluated based on diffusion pseudotime-based similarity measure $d_{2}$.

### Supplementary References


1. Wolfel R, Corman VM, Guggemos W, et al. Virological assessment of hospitalized patients with COVID-2019. *Nature*. May 2020;581(7809):465-469. doi:10.1038/s41586-020-2196-x
2. Adler JM, Martin Vidal R, Voss A, et al. A non-transmissible live attenuated SARS-CoV-2 vaccine. *Mol Ther*. Aug 2 2023;31(8):2391-2407. doi:10.1016/j.ymthe.2023.05.004
3. Nouailles G, Wyler E, Pennitz P, et al. Temporal omics analysis in Syrian hamsters unravel cellular effector responses to moderate COVID-19. *Nat Commun*. Aug 11 2021;12(1):4869. doi:10.1038/s41467-021-25030-7
4. Osterrieder N, Bertzbach LD, Dietert K, et al. Age-Dependent Progression of SARS-CoV-2 Infection in Syrian Hamsters. *Viruses*. Jul 20 2020;12(7)doi:10.3390/v12070779
5. Peidli S, Nouailles G, Wyler E, et al. Single-cell-resolved interspecies comparison identifies a shared inflammatory axis and a dominant neutrophil-endothelial program in severe COVID-19. *bioRxiv*. 2023:2023.08.25.551434. doi:10.1101/2023.08.25.551434
6. Trimpert J, Dietert K, Firsching TC, et al. Development of safe and highly protective live-attenuated SARS-CoV-2 vaccine candidates by genome recoding. *Cell Rep*. Aug 3 2021;36(5):109493. doi:10.1016/j.celrep.2021.109493
7. Andreotti S, Altmuller J, Quedenau C, et al. De Novo-Whole Genome Assembly of the Roborovski Dwarf Hamster (Phodopus roborovskii) Genome: An Animal Model for Severe/Critical COVID-19. *Genome Biol Evol*. Jul 2 2022;14(7)doi:10.1093/gbe/evac100
8. Stuart T, Butler A, Hoffman P, et al. Comprehensive Integration of Single-Cell Data. *Cell*. Jun 13 2019;177(7):1888-1902 e21. doi:10.1016/j.cell.2019.05.031
9. Schulte-Schrepping J, Reusch N, Paclik D, et al. Severe COVID-19 Is Marked by a Dysregulated Myeloid Cell Compartment. *Cell*. Sep 17 2020;182(6):1419-1440 e23. doi:10.1016/j.cell.2020.08.001
10. Patel NG, Bhasin A, Feinglass JM, Angarone MP, Cohen ER, Barsuk JH. Mortality, critical illness, and mechanical ventilation among hospitalized patients with COVID-19 on therapeutic anticoagulants. *Thrombosis Update*. 2021/01/01/ 2021;2:100027. doi:https://doi.org/10.1016/j.tru.2020.100027
11. Lotfollahi M, Wolf FA, Theis FJ. scGen predicts single-cell perturbation responses. *Nat Methods*. Aug 2019;16(8):715-721. doi:10.1038/s41592-019-0494-8
12. Kingma DP, Welling M. An Introduction to Variational Autoencoders. *Foundations and Trends® in Machine Learning*. 2019;12(4):307-392. doi:10.1561/2200000056
13. Wolf FA, Angerer P, Theis FJ. SCANPY: large-scale single-cell gene expression data analysis. *Genome Biol*. Feb 6 2018;19(1):15. doi:10.1186/s13059-017-1382-0
14. St»hle L, Wold S. Multivariate analysis of variance (MANOVA). *Chemometrics and Intelligent Laboratory Systems*. 1990/09/01/ 1990;9(2):127-141. doi:https://doi.org/10.1016/0169-7439(90)80094-M
15. Seabold S, Perktold J. Statsmodels: Econometric and Statistical Modeling with Python. 2010:
16. Haghverdi L, Buttner M, Wolf FA, Buettner F, Theis FJ. Diffusion pseudotime robustly reconstructs lineage branching. *Nat Methods*. Oct 2016;13(10):845-8. doi:10.1038/nmeth.3971
17. Lopez R, Regier J, Cole MB, Jordan MI, Yosef N. Deep generative modeling for single-cell transcriptomics. *Nat Methods*. Dec 2018;15(12):1053-1058. doi:10.1038/s41592-018-0229-2
18. Harris CR, Millman KJ, van der Walt SJ, et al. Array programming with NumPy. *Nature*. Sep 2020;585(7825):357-362. doi:10.1038/s41586-020-2649-2
19. Waskom M. seaborn: statistical data visualization. *The Journal of Open Source Software*. April 01, 2021 2021;6:3021. doi:10.21105/joss.03021
20. team Tpd. pandas-dev/pandas: Pandas (v2.1.3). Zenodo; 2023.
21. Isaac V, Sergei R, Fabian JT, Philipp A, Wolf FA. anndata: Annotated data. *bioRxiv*. 2021:2021.12.16.473007. doi:10.1101/2021.12.16.473007
22. Paszke A, Gross S, Chintala S, et al. Automatic differentiation in PyTorch. 2017:

23. Virtanen P, Gommers R, Oliphant TE, et al. SciPy 1.0: fundamental algorithms for scientific computing in Python. *Nat Methods*. Mar 2020;17(3):261-272. doi:10.1038/s41592-019-0686-2
24. Hunter JD. Matplotlib: A 2D Graphics Environment. *Computing in Science & Engineering*. 2007;9(3):90-95. doi:10.1109/MCSE.2007.55
25. Lun ATL, Richard AC, Marioni JC. Testing for differential abundance in mass cytometry data. *Nat Methods*. Jul 2017;14(7):707-709. doi:10.1038/nmeth.4295
26. Law CW, Chen Y, Shi W, Smyth GK. voom: Precision weights unlock linear model analysis tools for RNA-seq read counts. *Genome Biol*. Feb 3 2014;15(2):R29. doi:10.1186/gb-2014-15-2-r29
27. Aschenbrenner AC, Mouktaroudi M, Kramer B, et al. Disease severity-specific neutrophil signatures in blood transcriptomes stratify COVID-19 patients. *Genome Med*. Jan 13 2021;13(1):7. doi:10.1186/s13073-020-00823-5

### Supplemental Figures

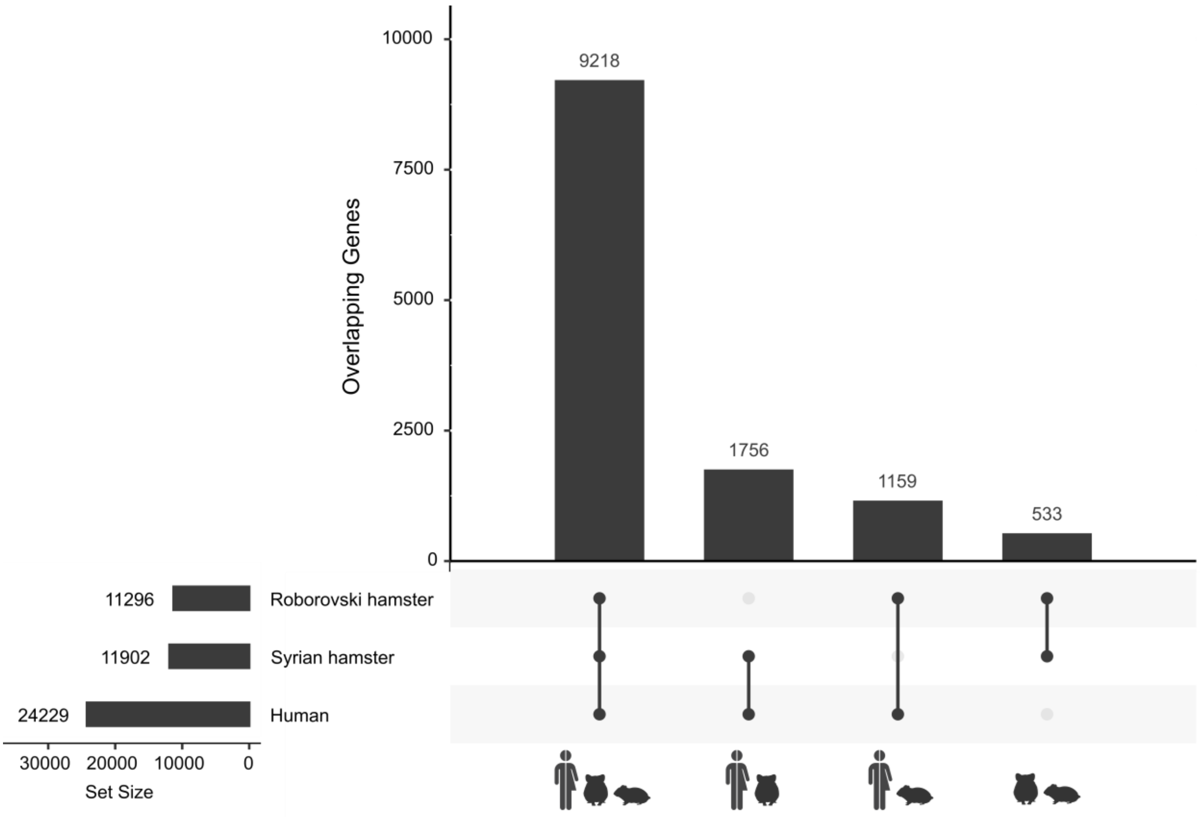


**Figure S1. Overlapping genes of interspecies datasets.** For the Roborovski hamster genes, human orthologues from genome assembly PHOROB[7] were used. The number to the left labelled as "Set Size" corresponds to the number of unique transcripts in each dataset. The number of genes found in multiple species is indicated in the graph, with the species combinations indicated by the connected dots on the x-axis, e.g. 9218 unique transcripts are found in all species.

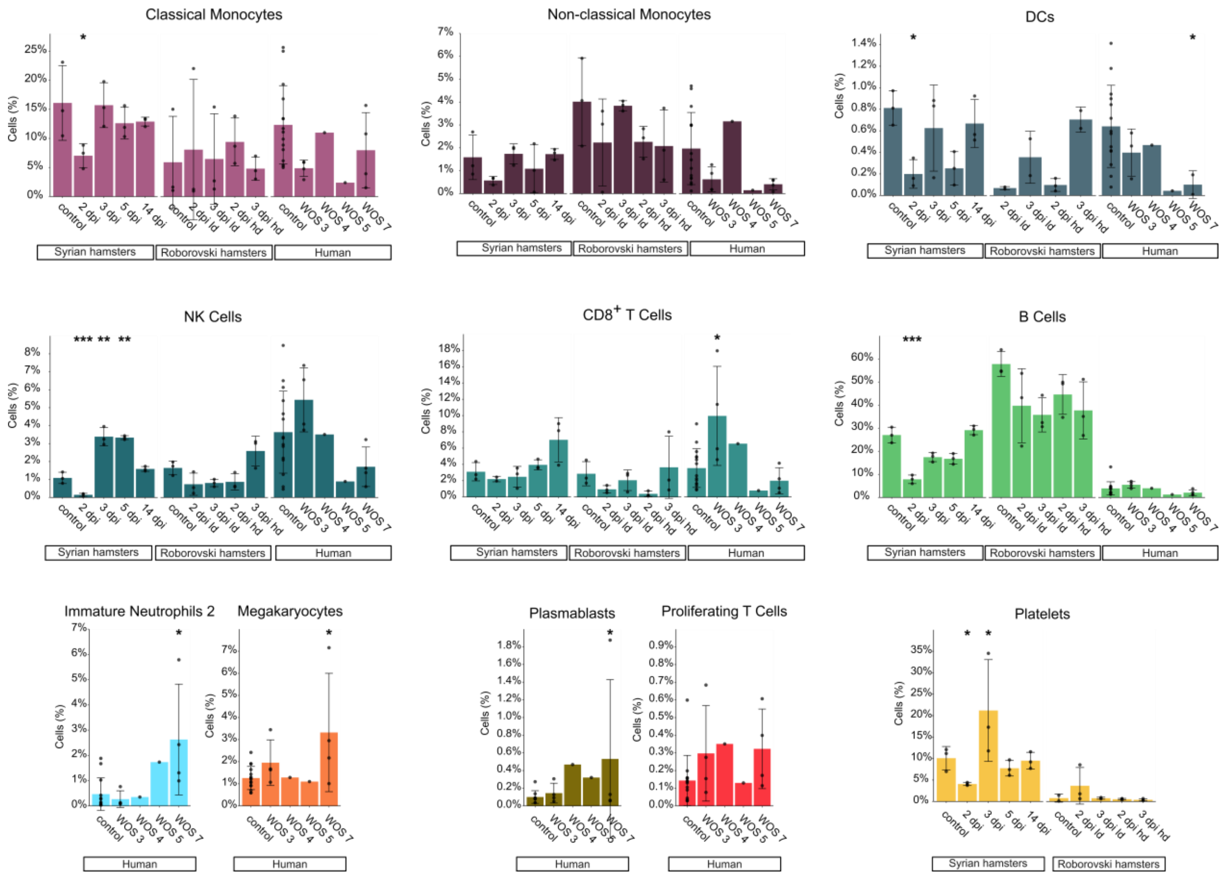


**Figure S2. Cellular recruitment in hamsters and humans.** (A) Barplots indicating frequency shifts of specific cell types at different severity levels in humans and time after infection in hamsters. Data display means ± SD. n = 3 animals per time point for hamsters. Human Data: Cohort 2 from Schulte-Schrepping, *et al*.[9]. To quantify significance of differential abundance of cell types, we applied a negative binomial generalized linear model as previously described[25]. * FDR ≤ 0.05, ** FDR ≤ 0.01, *** FDR ≤ 0.001. SD: standard deviation; FDR: global false discovery rate; WOS: WHO ordinal scale; NK-cells: natural killer cells; DCs: dendritic cells; dpi: days post infection; ld: low dose; hd: high dose.

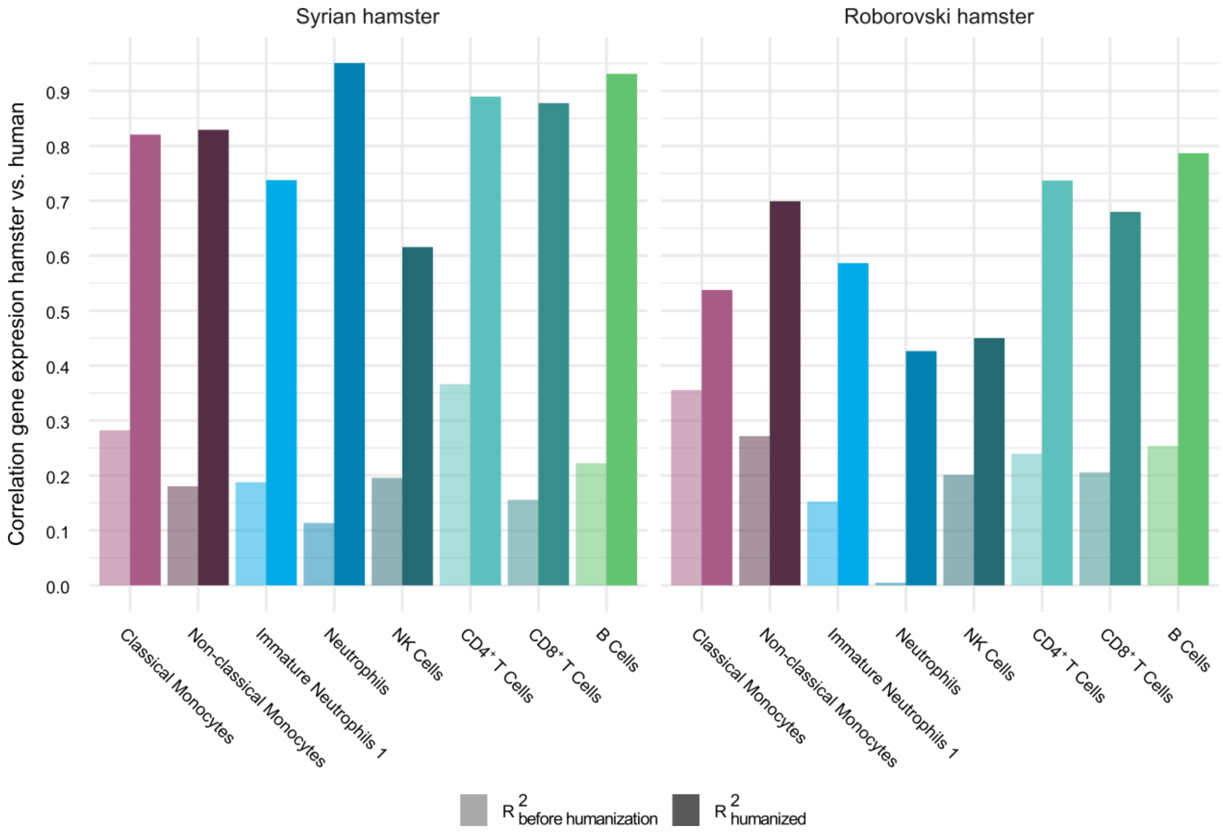


**Figure S3. Correlation measure of humanizing hamster cells using VAE neural network pipeline.** Barplots per cell type displaying $R^2$ Score of Spearman's rank correlation between hamster scRNA-seq data with human data before ("$R^2_{before\ humanization}$", light bars) and after ("$R^2_{humanized}$", bold bars) VAE-based humanization. Higher correlation with human data after humanization is indicative for successful VAE-based humanization. All Spearman's rank correlations showed positive values with the exception of $R^2_{before\ humanization}$ for Roborovski-hamster neutrophils. Across all tested cell types, the $R^2_{humanized}$ score displayed improved correlations. VAE: variational autoencoder; vs: versus; NK-cells: natural killer cells.

A

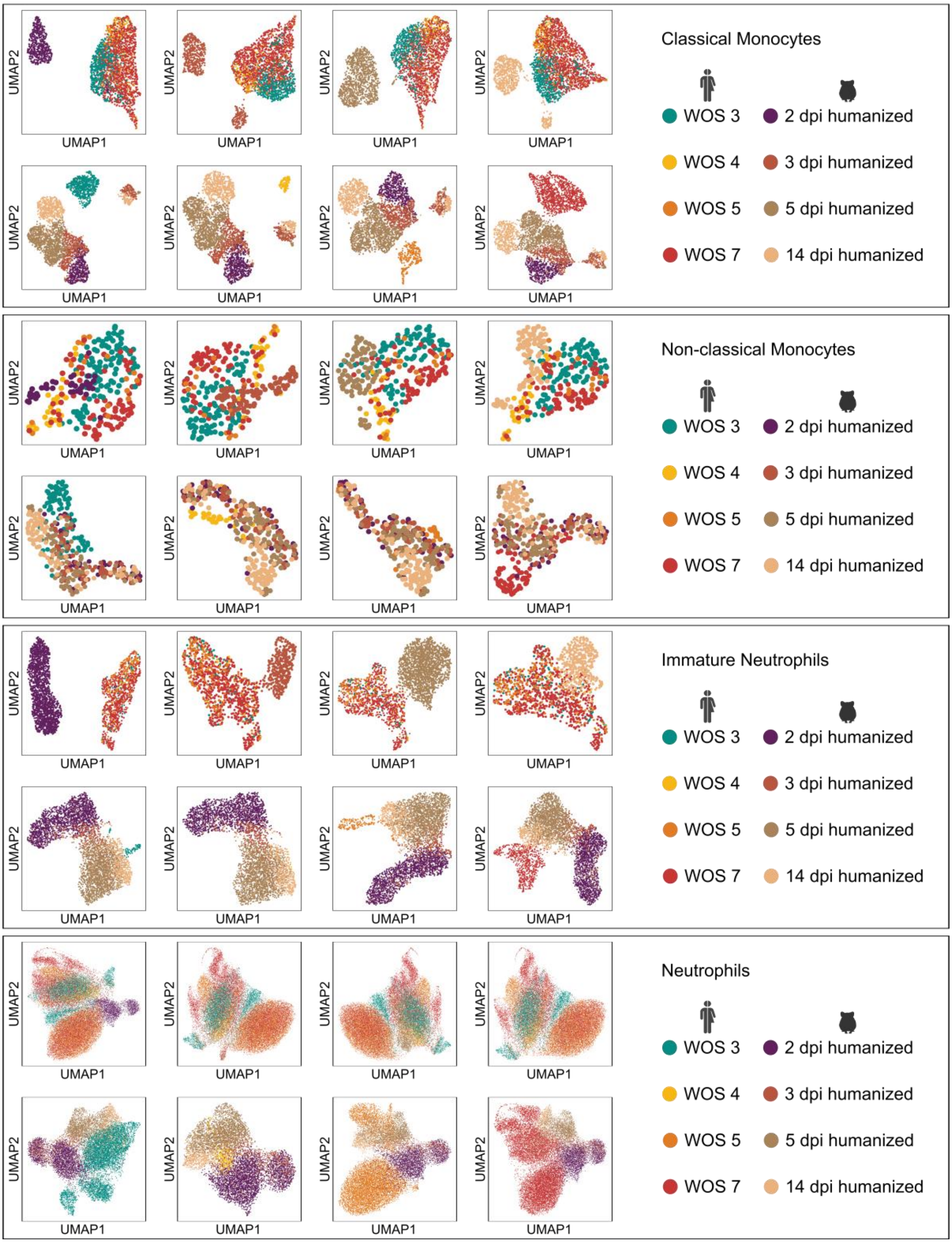
Classical Monocytes
WOS 3
WOS 4
WOS 5
WOS 7
2 dpi humanized
3 dpi humanized
5 dpi humanized
14 dpi humanized
Non-classical Monocytes
Immature Neutrophils
Neutrophils
UMAP1
UMAP2

**B**

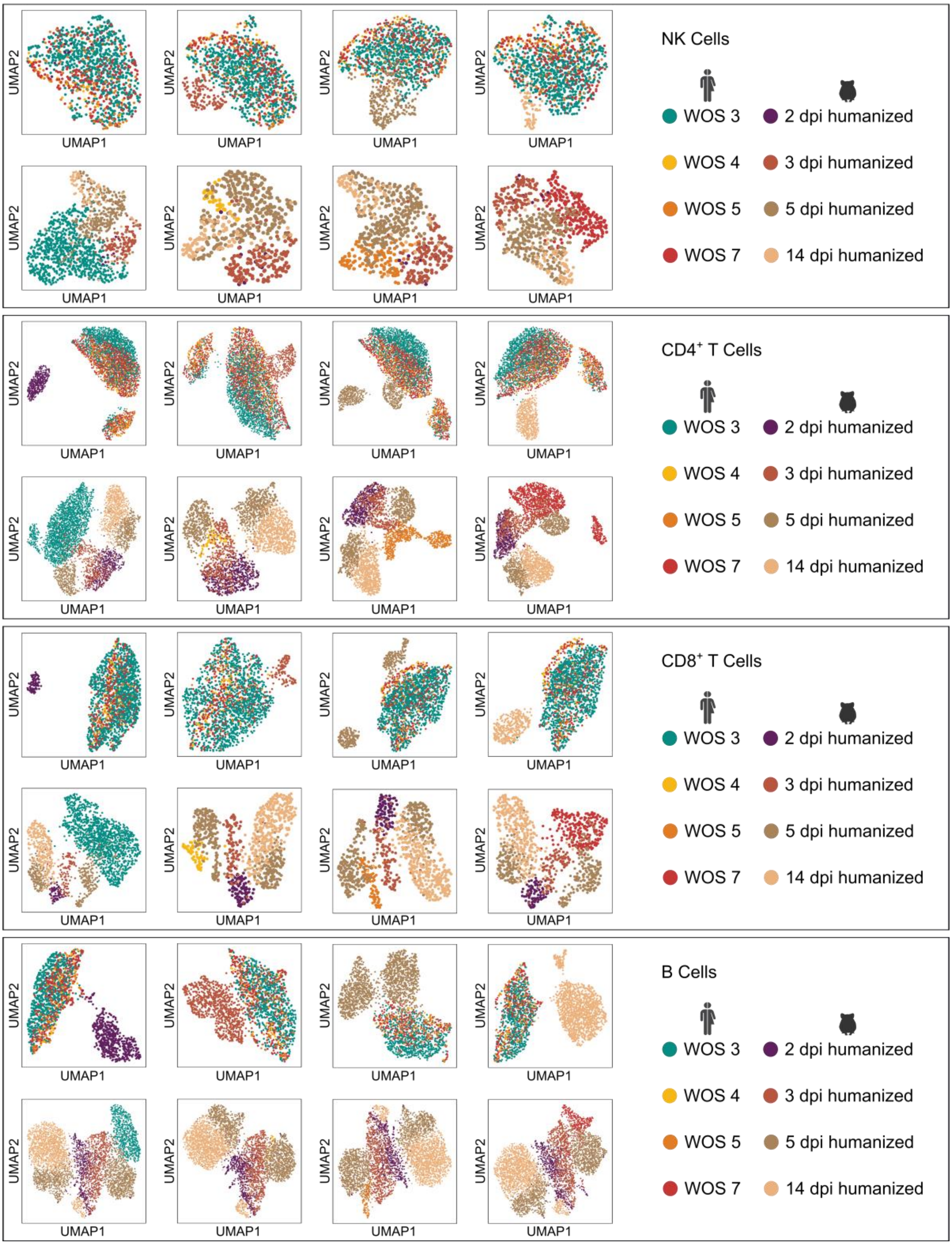
NK Cells
CD4+ T Cells
CD8+ T Cells
B Cells
WOS 3
WOS 4
WOS 5
WOS 7
2 dpi humanized
3 dpi humanized
5 dpi humanized
14 dpi humanized
UMAP1
UMAP2

C

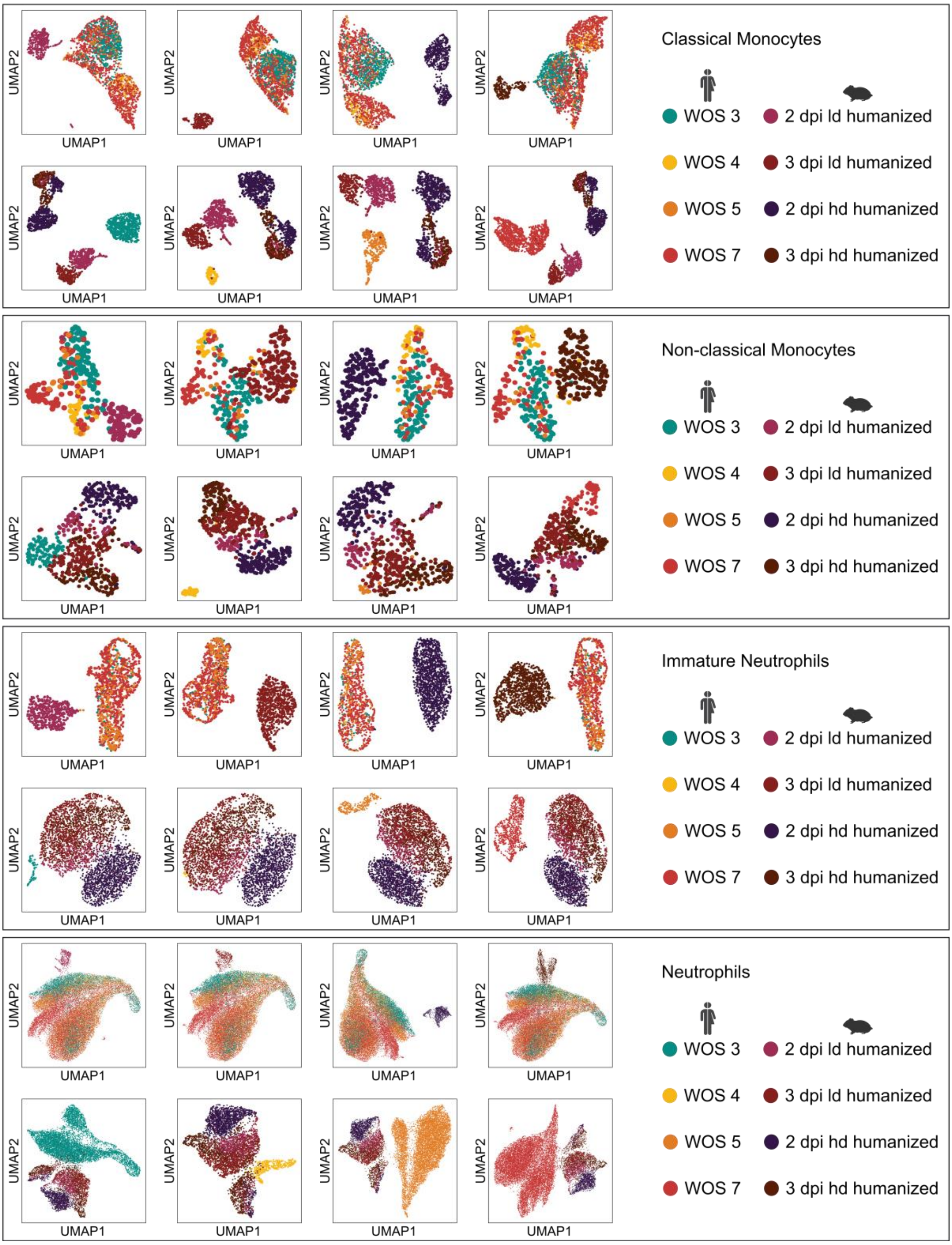
UMAP2
UMAP1
Classical Monocytes
WOS 3
WOS 4
WOS 5
WOS 7
2 dpi ld humanized
3 dpi ld humanized
2 dpi hd humanized
3 dpi hd humanized
Non-classical Monocytes
WOS 3
WOS 4
WOS 5
WOS 7
2 dpi ld humanized
3 dpi ld humanized
2 dpi hd humanized
3 dpi hd humanized
Immature Neutrophils
WOS 3
WOS 4
WOS 5
WOS 7
2 dpi ld humanized
3 dpi ld humanized
2 dpi hd humanized
3 dpi hd humanized
Neutrophils
WOS 3
WOS 4
WOS 5
WOS 7
2 dpi ld humanized
3 dpi ld humanized
2 dpi hd humanized
3 dpi hd humanized

D

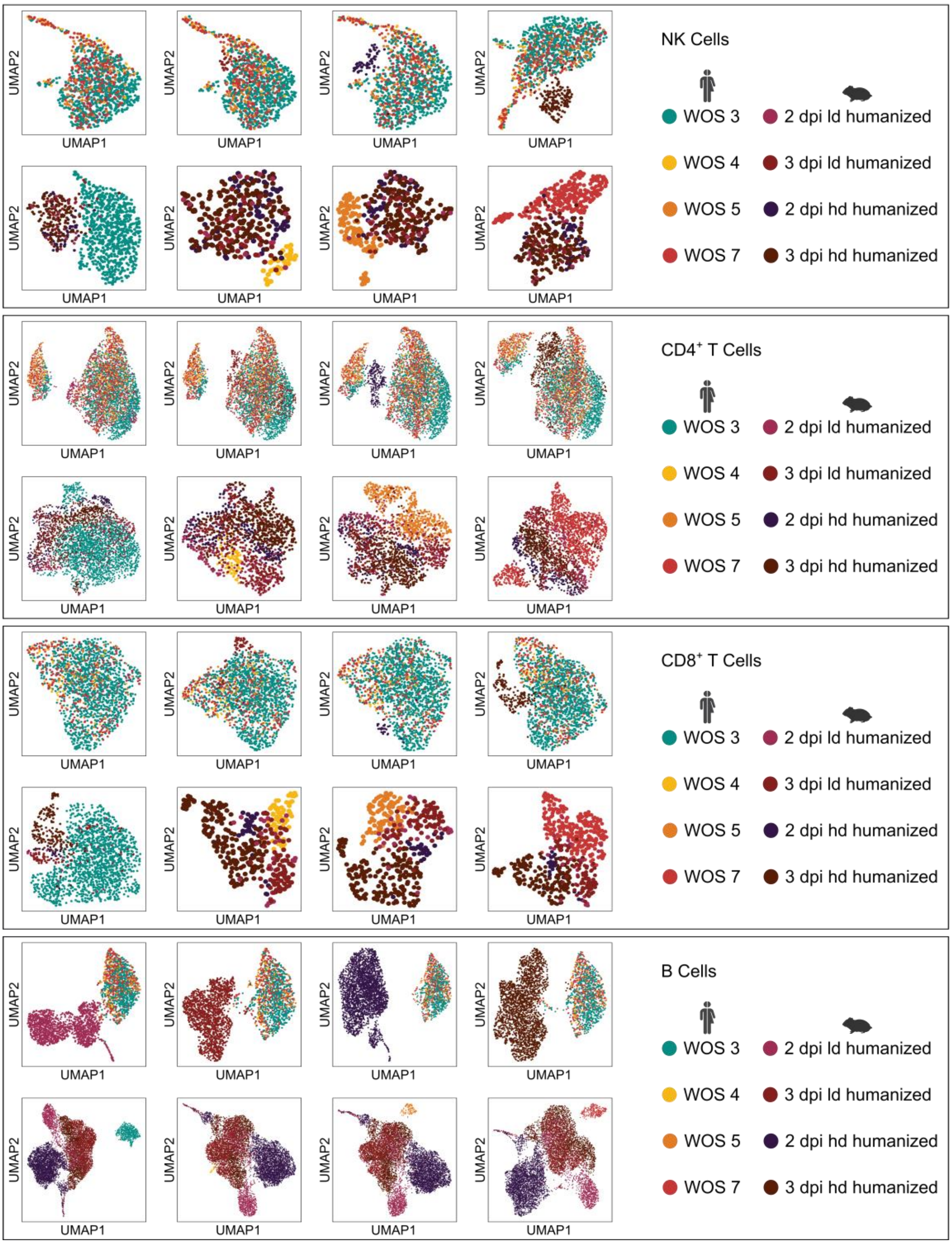


**Figure S4. Visualization of humanized hamster and human disease states within VAE neural network pipeline after application of species-shift-vector.** UMAP representations of VAE latent space embeddings after humanization of hamster disease states. Each UMAP plot shows one humanized hamster disease state jointly embedded with all human disease states (upper panels) or one human disease state jointly embedded with all humanized hamster disease states (lower panels) at the cell type level. (A) Syrian hamster and human classical monocytes, non-classical monocytes, immature neutrophils and neutrophils. (B) Syrian hamster and human NK-cells, $CD4^{+}$-T-cells, $CD8^{+}$-T cells and B-cells. (C) Roborovski hamster and human classical monocytes, non-classical monocytes, immature neutrophils and neutrophils. (D) Roborovski hamster and human NK-cells, $CD4^{+}$-T-cells, $CD8^{+}$-T-cells and B-cells. WOS: WHO ordinal scale; UMAP: uniform manifold approximation and projection for dimension reduction; VAE: variational autoencoder; NK-cells: natural killer cells; dpi: days post infection; ld: low dose; hd: high dose.

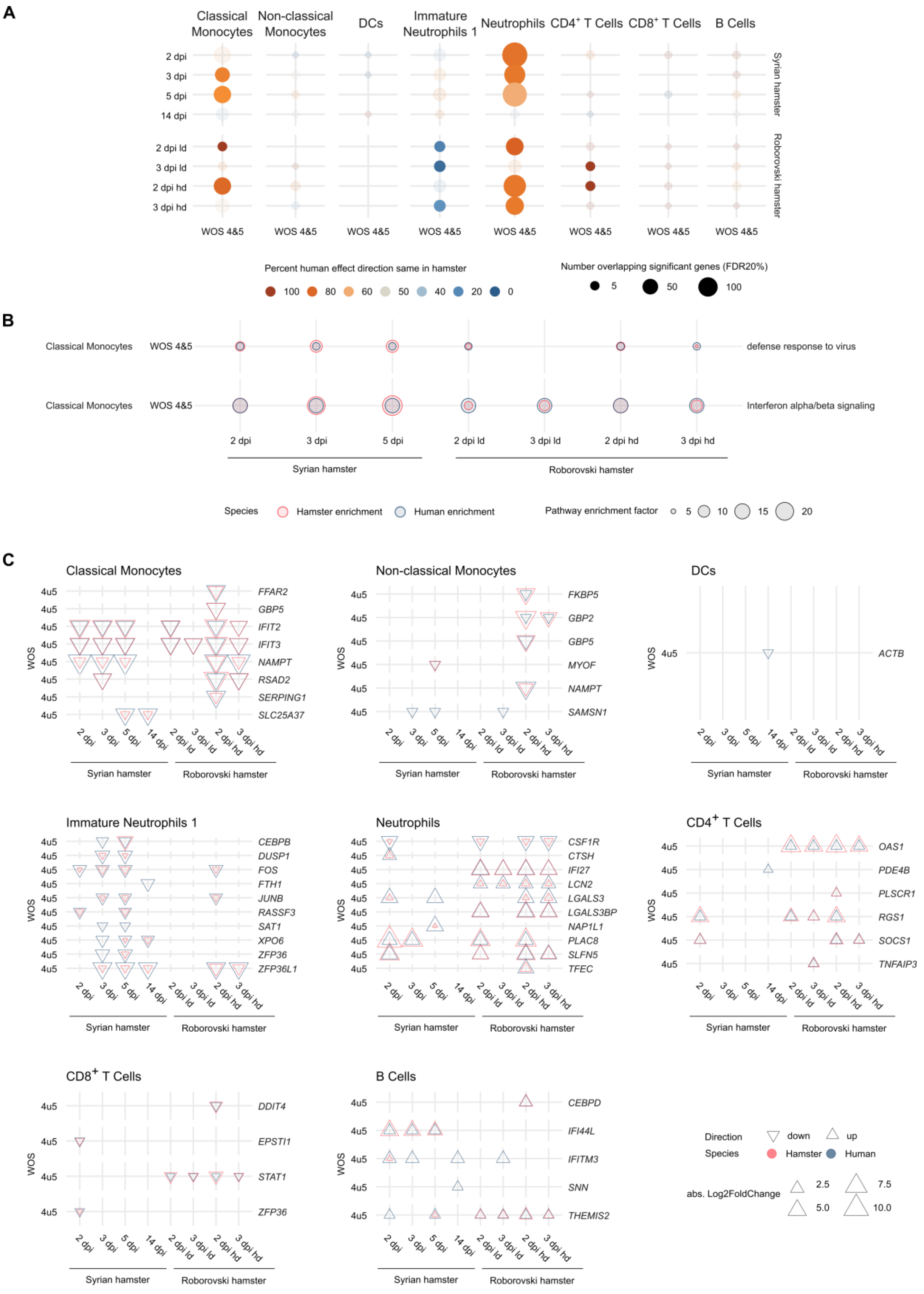


**Figure S5. Differential gene expression correlation of human and hamster disease states.** (A) Dotplot indicating significantly regulated genes that were regulated in the same direction in humans and hamsters. Shown in bright colors are conditions, in which significantly more (orange) or less (blue) than 50% of the overlapping genes were regulated in the same direction. (B) Dotplot of pathways of differentially expressed genes that are enriched in human and hamster species. (C) Dotplot displaying the top 10 genes for each cell type (ordered by absolute effect size at FDR 20%) that are regulated in corresponding direction (abs. Log2FoldChange) in humans and hamsters ordered by absolute effect size. Larger symbols correspond to larger effect sizes, direction of the triangle indicates up- or downregulation and color indicates the species. WOS: WHO ordinal scale; DCs: dendritic cells; dpi: days post infection; ld: low dose; hd: high dose. Differential expression analyses were performed in the Limma-Voom[26] framework. Multiple testing adjustments were applied using the false-discovery-rate according to Benjamini and Hochberg. To correlate fold-changes, all genes significant at false discovery rate (FDR) ≤20% in both species were used. Pathway enrichment analyses for globally significant pathways (*P* corrected ≤ 0.05) were performed using gprofiler2. Pathway enrichment factor is the ratio between observed and expected proportion of significant genes in each pathway.

A

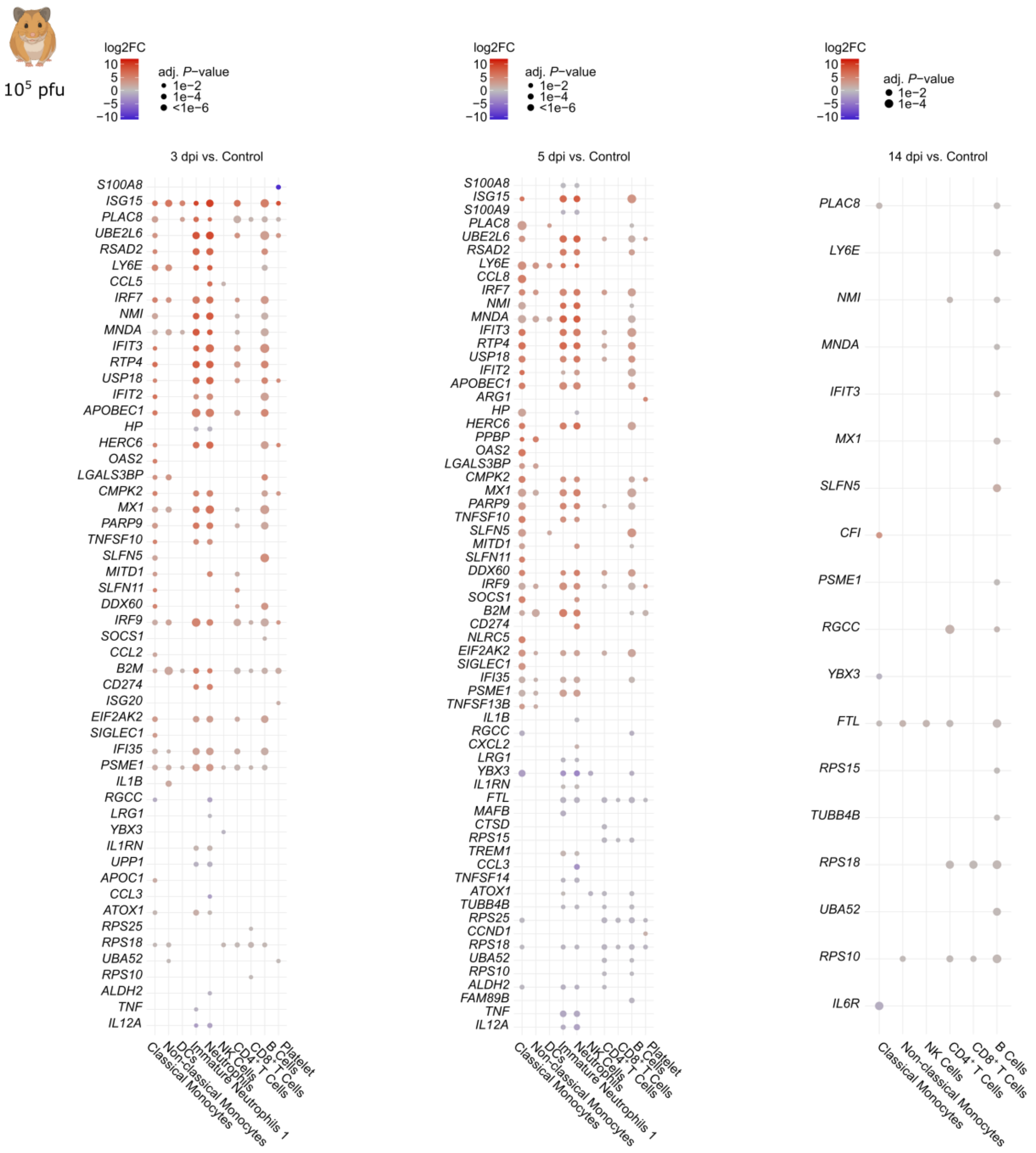

$10^5$ pfu
log2FC
10
5
0
−5
−10
adj. P−value
1e−2
1e−4
<1e−6
3 dpi vs. Control
S100A8
ISG15
PLAC8
UBE2L6
RSAD2
LY6E
CCL5
IRF7
NMI
MNDA
IFIT3
RTP4
USP18
IFIT2
APOBEC1
HP
HERC6
OAS2
LGALS3BP
CMPK2
MX1
PARP9
TNFSF10
SLFN5
MITD1
SLFN11
DDX60
IRF9
SOCS1
CCL2
B2M
CD274
ISG20
EIF2AK2
SIGLEC1
IFI35
PSME1
IL1B
RGCC
LRG1
YBX3
IL1RN
UPP1
APOC1
CCL3
ATOX1
RPS25
RPS18
UBA52
RPS10
ALDH2
TNF
IL12A
Classical Monocytes
Non-classical Monocytes
DCs
Immature Neutrophils 1
Neutrophils
NK Cells
CD4+ T Cells
CD8+ T Cells
B Cells
Platelet
5 dpi vs. Control
S100A8
ISG15
S100A9
PLAC8
UBE2L6
RSAD2
LY6E
CCL8
IRF7
NMI
MNDA
IFIT3
RTP4
USP18
IFIT2
APOBEC1
ARG1
HP
HERC6
PPBP
OAS2
LGALS3BP
CMPK2
MX1
PARP9
TNFSF10
SLFN5
MITD1
SLFN11
DDX60
IRF9
SOCS1
B2M
CD274
NLRC5
EIF2AK2
SIGLEC1
IFI35
PSME1
TNFSF13B
IL1B
RGCC
CXCL2
LRG1
YBX3
IL1RN
FTL
MAFB
CTSD
RPS15
TREM1
CCL3
TNFSF14
ATOX1
TUBB4B
RPS25
CCND1
RPS18
UBA52
RPS10
ALDH2
FAM89B
TNF
IL12A
Classical Monocytes
Non-classical Monocytes
DCs
Immature Neutrophils 1
Neutrophils
NK Cells
CD4+ T Cells
CD8+ T Cells
B Cells
Platelet
14 dpi vs. Control
adj. P−value
1e−2
1e−4
PLAC8
LY6E
NMI
MNDA
IFIT3
MX1
SLFN5
CFI
PSME1
RGCC
YBX3
FTL
RPS15
TUBB4B
RPS18
UBA52
RPS10
IL6R
Classical Monocytes
Non-classical Monocytes
NK Cells
CD4+ T Cells
CD8+ T Cells
B Cells

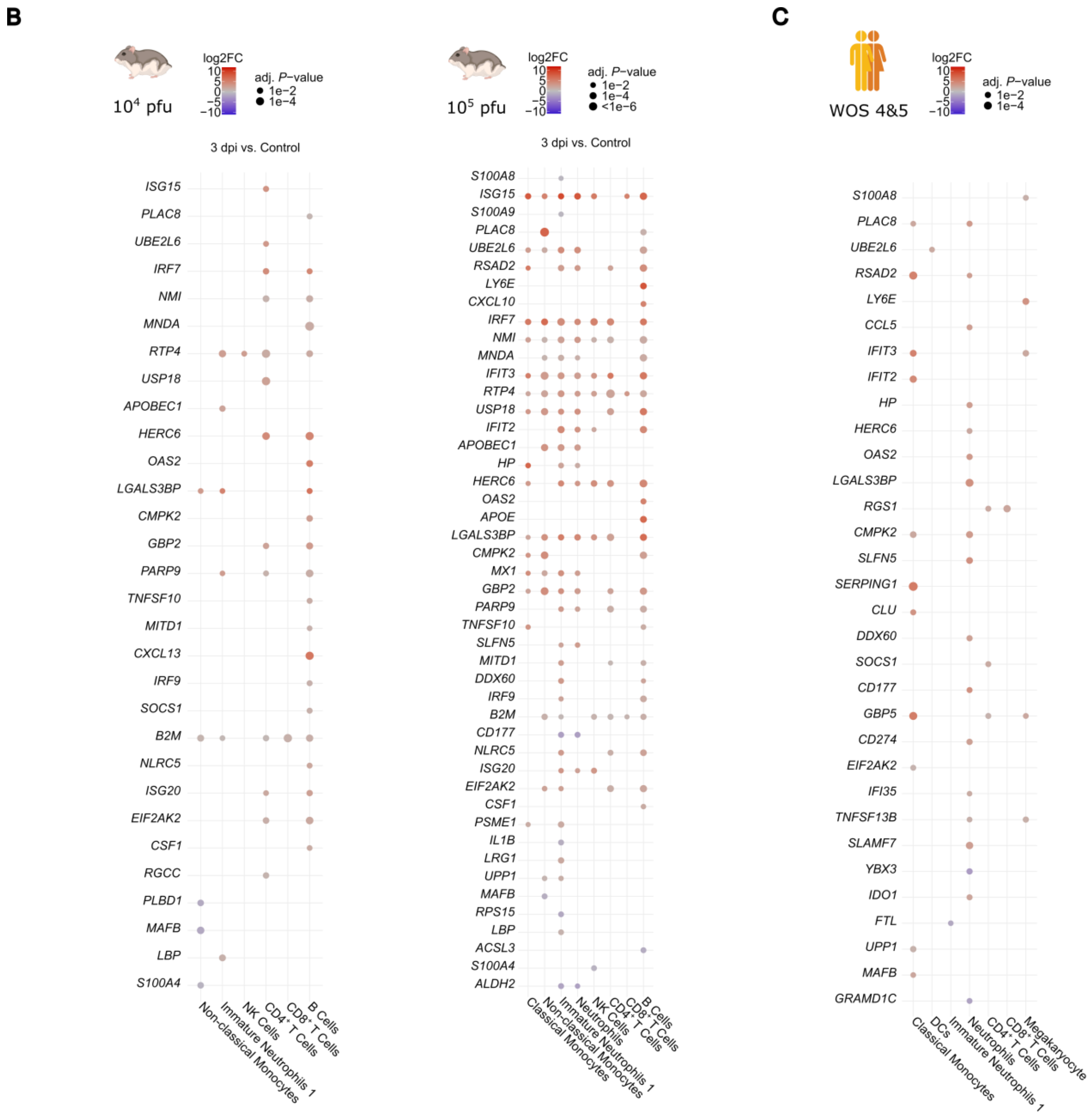


**Figure S6. COVID-19 mediators and gene expression patterns amongst relevant cell types.** Dotplots displaying significantly up- or downregulated genes linked to inflammation in (A) Syrian hamsters by days post-infection compared to controls, (B) Roborovski hamsters by days post-infection compared to controls, (C) COVID-19 patients by WOS severity level compared to controls. NK cells: natural killer cells; DCs: dendritic cells; WOS: WHO ordinal scale; dpi: days post infection; pfu: plaque-forming units; log2FC: log2FoldChange. Differential expression analyses were performed in the Limma-Voom[26] framework. Adj. *P*-value: adjusted *P*-values were calculated according to Benjamini and Hochberg.

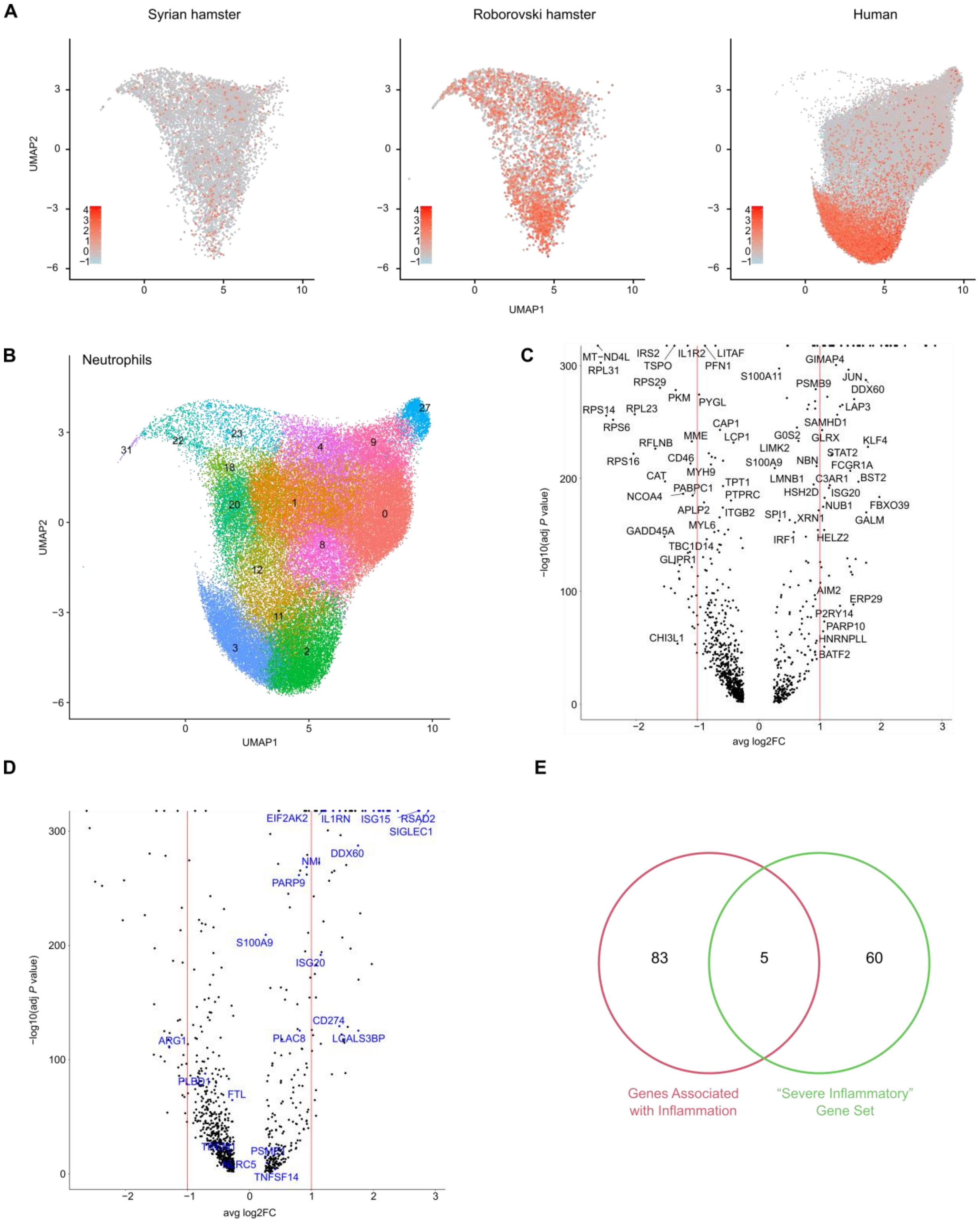

**Figure S7. Highly inflammatory response triggered by COVID-19 in human neutrophils is limited to a subset.** (A) UMAP plots displaying expression of the previously described “severe inflammatory” gene set[27] amongst peripheral blood neutrophils in humans or indicated hamster species. (B) UMAP plot displaying the subclustering of neutrophils. (C) Volcano plot displaying genes differentially expressed between neutrophil clusters 2 and 3 of WOS 5 and WOS 7 patients compared to all other neutrophil clusters of these patients. (D) Volcano plot indicating differential expression of genes associated with inflammation. (C, D) Only significantly differentially regulated genes (adj *P*-values ≤ 0.05 after Bonferroni correction for multiple testing) are shown. (E) Venn diagram displaying numbers of overlapping genes expressed in the merged interspecies dataset between the “severe inflammatory” gene set[27] (green) and genes associated with inflammation (red). WOS: WHO ordinal scale; UMAP: uniform manifold approximation and projection for dimension reduction; adj: adjusted.
